# Experimental manipulation of selfish genetic elements links genes to microbial community function

**DOI:** 10.1101/608752

**Authors:** Steven D. Quistad, Guilhem Doulcier, Paul B. Rainey

## Abstract

Microbial communities underpin earth’s biological and geochemical processes, but their complexity hampers understanding. Motivated by the challenge of diversity and the need to forge ways of capturing dynamical behaviour connecting genes to function, biologically independent experimental communities comprising hundreds of microbial genera were established from garden compost and propagated on nitrogen-limited minimal medium with cellulose (paper) as sole carbon source. After one year of bi-weekly transfer, communities retained hundreds of genera. To connect genes to function we used a simple experimental manipulation that involved periodic collection of selfish genetic elements (SGEs) from separate communities, followed by pooling and redistribution across communities. The treatment was predicted to promote amplification and dissemination of SGEs and thus horizontal gene transfer (HGT). Confirmation came from comparative metagenomics, which showed substantive movement of ecologically significant genes whose dynamic across space and time could be followed. Enrichment of genes implicated in nitrogen metabolism, and particularly ammonification, prompted biochemical assays that revealed a measurable impact on community function. Our simple experimental strategy offers a conceptually new approach for unravelling dynamical processes affecting microbial community function.

## Introduction

Microbial communities underpin all major biological and biogeochemical processes [1]. Appropriate functioning of communities has direct implications for health of people [2, 3], terrestrial [4, 5], marine [6] and freshwater [7] ecosystems and even global climate [8]. While the species composition of many communities have been exhaustively documented [9, 10], the link between genes and community function is poorly understood [1].

New strategies for investigation are required that are process-focused [11]. Ideally such approaches will provide knowledge on the complex interconnections between community members, their combined effects, including feedbacks that shape the evolution of community members [12–15]. Recent advances draw upon new strategies for linking patterns of sequence diversity in metagenomic data sets to population genetic processes [16–19]. Knowledge of such processes provides a link between variation and the likelihood that particular traits fix.

Equally desirable are approaches that provide a link between genes, their spatial and temporal dynamics, and community function, achieved using strategies that leave as far as possible the natural complexity of communities undisturbed. Here we show — in a proof-of-principle experiment — that this can be achieved via an experimental strategy that combines theory governing the behaviour of selfish genetic elements [20–23] (SGEs) and expected effects on horizontal gene transfer (HGT), with approaches from experimental evolution [24, 25], comparative metagenomics [26] and functional assay.

## Results

### Diversity in experimental communities

Experimental communities were established from a diverse primary source: garden compost. Ten independent 1 g samples of compost were taken from ten different regions of a single 1 m^3^ compost heap and each placed in one of ten 140 ml bottles (mesocosms) containing 20 ml nitrogen-limited minimal M9 medium plus cellulose (a 4 cm^2^ piece of paper) as sole carbon source. Mesocosms were incubated on a laboratory bench without shaking and with lids left unsealed. After a two week period the initial piece of paper from each mesocosm was separately transferred to a new bottle containing fresh M9 medium plus a new piece of paper. Following a further two week incubation period we considered the communities to have adjusted to laboratory conditions. At this time point mesocosms were vortex-mixed and 1 ml of cellulosic slurry from each bottle was transferred to *two* new mesocosms containing 19 ml fresh M9 medium plus paper giving a total of 20 paired communities. This transfer, four weeks after initial establishment, was deemed time point zero (T_0_). Thereafter and for the ensuing 48 weeks, communities were serially propagated every two weeks (T_1_ to T_24_) by transferring 1 ml of slurry to fresh bottles. One set of each pair of bottles was labelled “*vertical*” (vertical communities (VC)) and the second of each pair labelled “*horizontal*” (horizontal communities (HC)). Details of the treatments and their significance are elaborated below, but at this stage, with focus solely on the question of diversity in cellulose-based mesocosms, we acknowledge treatment effects by name only.

Diversity through 48 weeks of propagation was addressed by metagenomics: total DNA was extracted from each of the ten T_0_, and twenty T_1_ and T_24_ communities and the data interrogated for sequence reads mapping to 16S rDNA (Supplementary Table 1). Communities at T_0_ harboured on average ~140 bacterial genera (SD ± 23). At T_24_ the number of genera detected increased in all communities to an average of ~200 (SD ± 32), indicating an increase in abundance of rare types (Supplementary Table 1, Supplementary Fig. 1a). Diversity levels were highly similar in both HCs and VCs (Supplementary Fig. 1a-c). Rank abundance distributions from pooled HCs and VCs at T1 and T24 are shown in Fig. 1 and reveal the tail of the T_24_ distribution to be markedly flatter in both cases, consistent with an overall tendency for rare types to have increased during the course of the year long selection. Rank abundance distributions for each T_1_ and T_24_ HC and VC, plus the rank order (and change therein) of the most common genera are shown in Supplementary Figs 2a and 2b. Supplementary Fig. 3 is a multidimensional scaling plot showing that communities at the start and end of the experiment differed markedly in genera composition and there was no sign of convergence after 48 weeks. Among the most abundant genera, only two harbour species commonly associated with cellulolytic ability [27] (*Cytophaga* and *Cellvibrio*), while others are better known for roles in ammonification [28] (*Azospirillum* and *Paenibacillus* (nitrogen fixation); *Rhodanobacter* and *Pseudomonas* (dissimilatory nitrate reduction / denitrification)) (Supplementary Figs 2a and 2b). Single celled eukaryotes were also present in all mesocosms with several unexpectedly maintaining populations of nematodes through the 48-week selection period.

**Figure 1:**
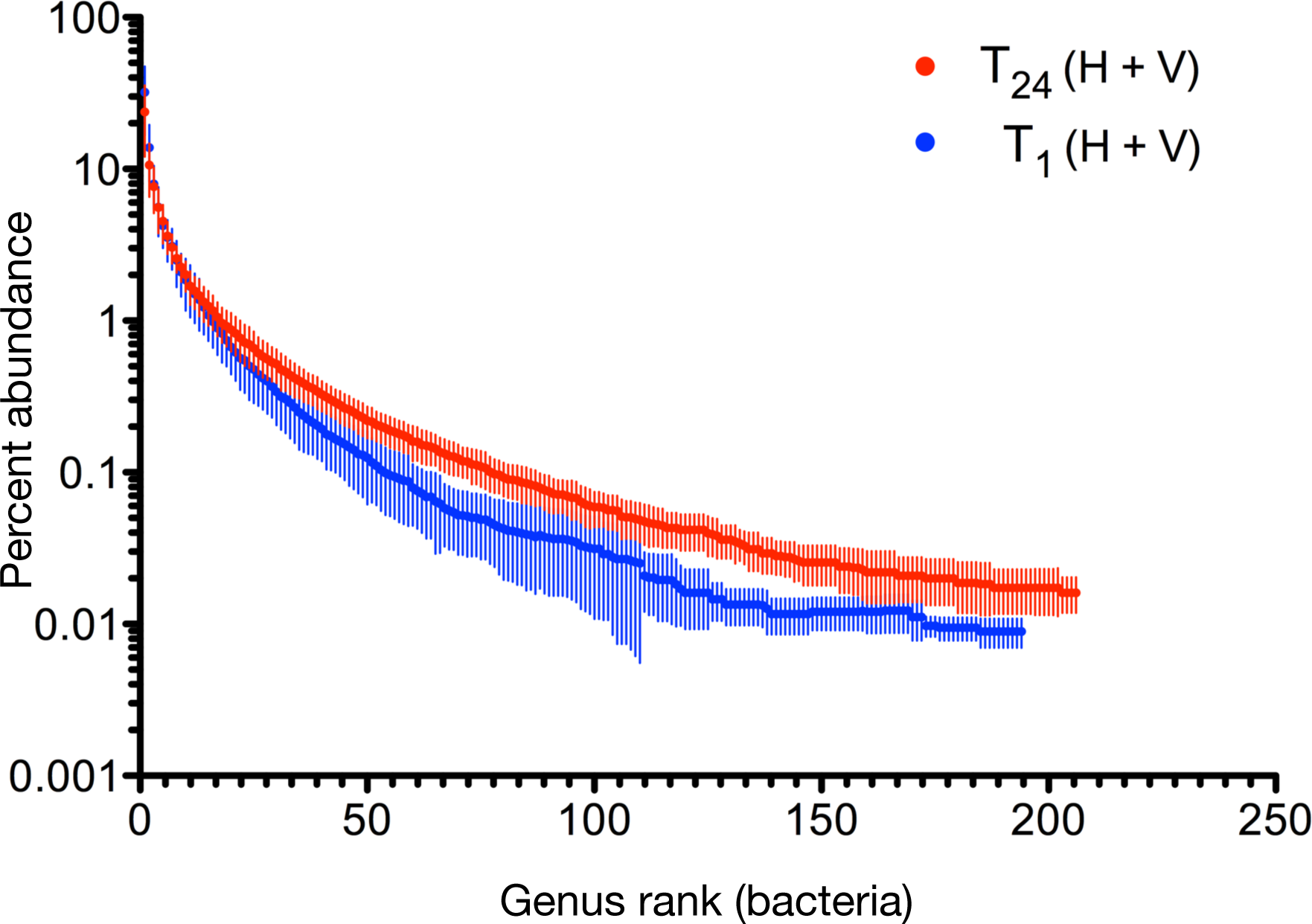
Genus-level rank abundance curves. The blue line depicts the genus-level rank abundance curve for bacteria at time point T_1_ two weeks after the divide of founding communities into horizontal and vertical treatments. The red line is at time point T_24_ (48 weeks). Because there were no differences between vertical and horizontal communities at T_1_ or T_24_ data from both regimes were combined. Data are means and standard deviations from 20 mesocosms.

### Manipulating community-level process

An important goal is development of strategies that link genes to community function by way of process. An ideal experiment is one that alters a single ecological or evolutionary process and compares the outcome to a set of replicates in which the process is unaffected. For example, evolutionary biologists interested in the evolutionary consequences of sex have taken advantage of yeast in which it is possible to genetically engineer sexual and asexual types that are otherwise isogenic [29, 30]. Differences in evolutionary outcome between sexual and asexual types after a period of selection under identical environmental conditions can thus be attributable to sex.

We reasoned that a similar manipulation might be possible in the context of microbial communities. Recombination mediated by horizontal gene transfer (HGT) is loosely analogous to sex, albeit to a form of sex that takes place at the level of communities. Moreover, like genetic manipulations affecting sexual reproduction in yeast, the extent of HGT can be controlled by manipulating the evolutionary fate of selfish genetic elements (SGEs) [20, 23]. Such elements, including phages, plasmids, transposons and integrative and conjugative elements are prime vehicles for the movement of ecologically significant genes among diverse bacteria [31–33]. We note that when SGEs acquire genes that aid host survival they are no longer “selfish” and are more appropriately referred to as “mobile” genetic elements, however, the theory we draw upon comes from thinking about elements that are costly to maintain and thus we continue to use the more generic SGE label.

The critical manipulation is one that affects the likelihood that SGEs encounter new hosts. Frequent exposure to new hosts drives dissemination and evolution of SGEs and linked genes [23]. Conversely, in the absence of new hosts, and thus opportunity for infectious spread, selection is powerless to prevent loss of transfer ability leading to degradation of SGEs and thus reduction in HGT. That loss occurs is evidenced by the remnants of prophages that litter bacterial genomes [34].

The *vertical* and *horizontal* treatment designations referred to above reflect manipulations that are predicted to either restrict, or facilitate, respectively, HGT mediated by SGEs. The vertical regime involved serial transfer of material from mesocosm to mesocosm. SGEs within these communities were denied opportunity to encounter new hosts: transfer of SGEs was vertical and the extent of HGT is expected to be limited (Fig. 2) (and eliminated altogether over longer timescales).

Communities subject to the horizontal regime were transferred by serial dilution to fresh mescosms as per the vertical regime, but received additionally — at the time of serial transfer — an aliquot of “SGE cocktail”. The SGE cocktail was a pooled sample of SGEs derived from each horizontal mesocosm, which was then redistributed among each HC (Fig. 2). SGEs within these communities were thus repeatedly (every two weeks) provided with opportunity to encounter new hosts. For example, phages from HC 1 could infect hosts present in HCs 2-9 and may additional mobilise ecologically significant genes acquired en route. The horizontal regime thus breathes evolutionary life into SGEs and HGT is expected to be rampant.

**Figure 2:**
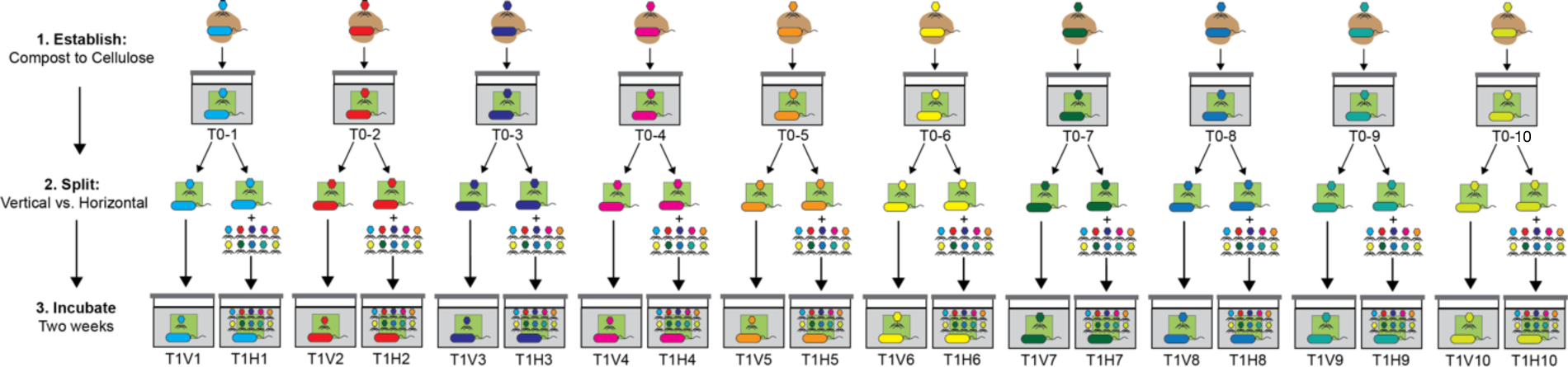
Experimental protocol for manipulation SGEs and thus the extent of HGT. Cellulose-degrading microbial communities were established by placing ten independent 1 gram samples of fresh compost (taken from separate locations in a single compost heap) into minimal M9 medium contained within a glass mesocosm with paper as the sole carbon source. Following four weeks of incubation at room temperature communities were homogenized and divided in two. Thereafter — and for the next 48 weeks — one of each pair was subject to a “vertical” and the other a “horizontal” transfer regime. The vertical regime involved serial transfer of a sample from each community every two weeks. The horizontal regime involved the same serial transfer protocol, but additionally, at the time of transfer a sample of supernatant was collected from each of the ten horizontal communities and passed through a 0.2 µm filter producing a “SGE-cocktail”. This sample of ten SGE-cocktails was pooled (indicated by the 10 different coloured phage cartoons) and redistributed across all horizontal microcosms. Note: DNA was not extracted from the SGE-cocktail prior to mixing (or subsequently).

### Detection of SGE activity

To determine whether the horizontal regime promoted movement of SGEs, total DNA was extracted from each T_0_ community and each of the 20 T_1_ communities (two weeks after the split of communities into vertical and horizontal treatments) and sequenced (Fig. 2 and Supplementary Table 2). Analysis was straightforward given the paired experimental design. For each set of T_0_ and descendant HCs and VCs (e.g., T_0_-1 and descendants T_1_V-1 and T_1_H-1) DNA sequence data were interrogated to identify reads found solely in (and therefore unique to) the horizontal mesocosm (e.g., T_1_H-1). Such unique reads are likely to stem from SGEs, or be associated with SGE activity, originating from an allopatric community.

Analysis of the sequences unique to each horizontal community resulted in identification of ~26 million unique reads from a total of ~152 million reads across all ten communities, with an average of ~2.6 million unique reads per community (Supplementary Table 3). Assembly of reads into contigs revealed an average of 3,352 contigs per HC greater than 1 kb in length. The mean maximum contig size per assembly was 82 kb (Supplementary Table 3).

After extracting Open Reading Frames (ORFs), interrogation of the Conserved Domain Database (CDD) [35] showed 1,279 ORFs predicted to encode phage-associated proteins involved in capsid formation, baseplate assembly, and phage-induced lysis of bacterial cells (Supplementary Table 3). An example of a contig (T_1_H-1_35969) containing numerous genes characteristic of phages is shown in Supplementary Fig. 4a.

The distribution and abundance of contig T_1_H-1_35969 and 19 other representative phage-like entities (Supplementary Table 4) across independent mesocosms was determined by mapping total reads from each horizontal and vertical metagenome onto each of the 20 phage-like contigs (Fig. 3). The predominance of these contigs in HCs is evidence of rapid amplification and dissemination of genetic material. T_1_H-1_35969, originally assembled from the unique set of sequences from T_1_H-1, was also present in both T_1_H-2 and T_1_V-2 and was therefore likely present in the founding community (T_0_-2). Within two weeks this contig had amplified and spread to horizontal communities 1, 5 and 6 (Fig. 3). Contig T1H2_39307 was below the level of detection in all vertical treatments, but two weeks after imposition of the horizontal treatment, this contig was detected in seven of the independent HCs (Fig. 3, Supplementary Table 4). From the community perspective, less than four of the 20 phage-like contigs were detected on average per VC, whereas the mean number of contigs per HCs was 11 (Fig. 3).

In addition to phage-like contigs, assemblies of the unique sequences from HCs contained genes predicted to encode traits of likely ecological significance (Supplementary Fig. 5). These include genes implicated in cellulose degradation, defence against SGE invasion, nitrogen metabolism and siderophore biosynthesis (Supplementary Table 5) [36–38]. Particularly notable was enrichment of reads mapping to *β*-glucosidase genes that define the rate-limiting step in cellulose degradation [39]. On occasion these ecologically significant genes were linked to genes encoding features found on SGEs (Supplementary Table 6), but often these contigs lacked such features (Supplementary Fig. 4b). Nonetheless they were amplified and disseminated through the course of the selection experiment with the same dynamic as expected of a mobile genetic element (see below).

**Figure 3:**
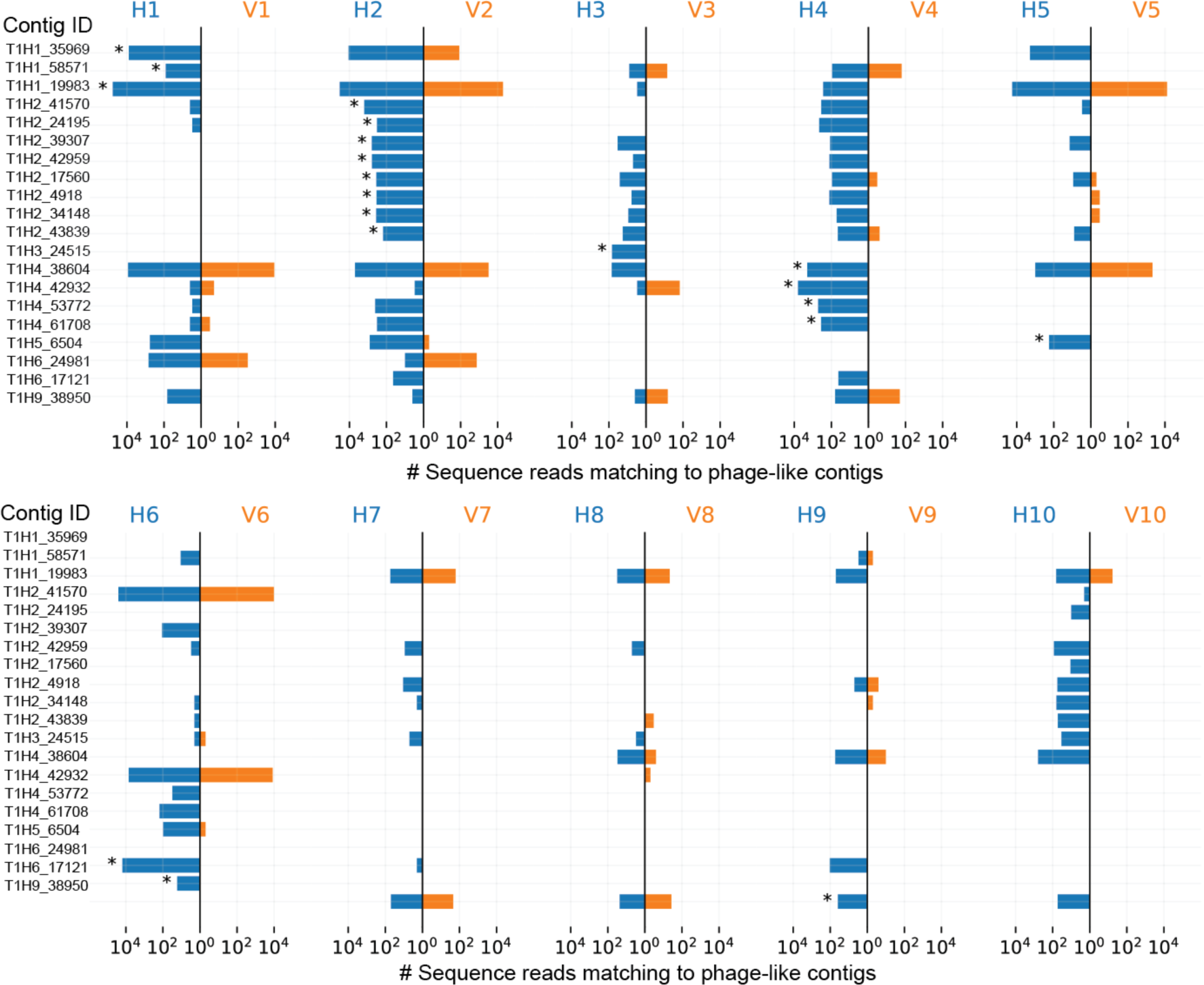
Movement of phage-like elements between horizontal communities. Twenty phage-like contigs assembled from sequence reads unique to horizontal communities are listed on the y-axes. Each vertical line represents a single pair of mesocosms with communities subject to the horizontal and vertical regimes depicted in blue and orange, respectively. The x-axis of the butterfly plot is the number of reads mapping to each contig. Astrices denote the source community from which the contigs were assembled.

### Tracking the dynamics of SGE amplification and dissemination

Given evidence indicative of the amplification and dissemination of DNA via the horizontal treatment just two weeks after splitting T_0_ communities into VCs and HCs, we complemented existing DNA sequence from T_0_ and T_1_ and T_24_ with DNA extracted from all HCs and VCs at time points T_2_, T_3_, T_4_, T_10_, T_16_, and T_20_ yielding a total of 180 metagenomes (5.2 billion reads, each ~150 bp). These 180 metagenomic data sets (nine time points, 20 communities per time point (Supplementary Table 2)) spread across 48 weeks allowed the abundance and distribution of SGEs and associate genes to be determined by read mapping to contigs assembled from T_0_ data. The spatial and temporal dynamics of two phage-like contigs and two contigs encoding *β*-glucosidases are shown in Fig. 4a-d.

**Figure 4:**
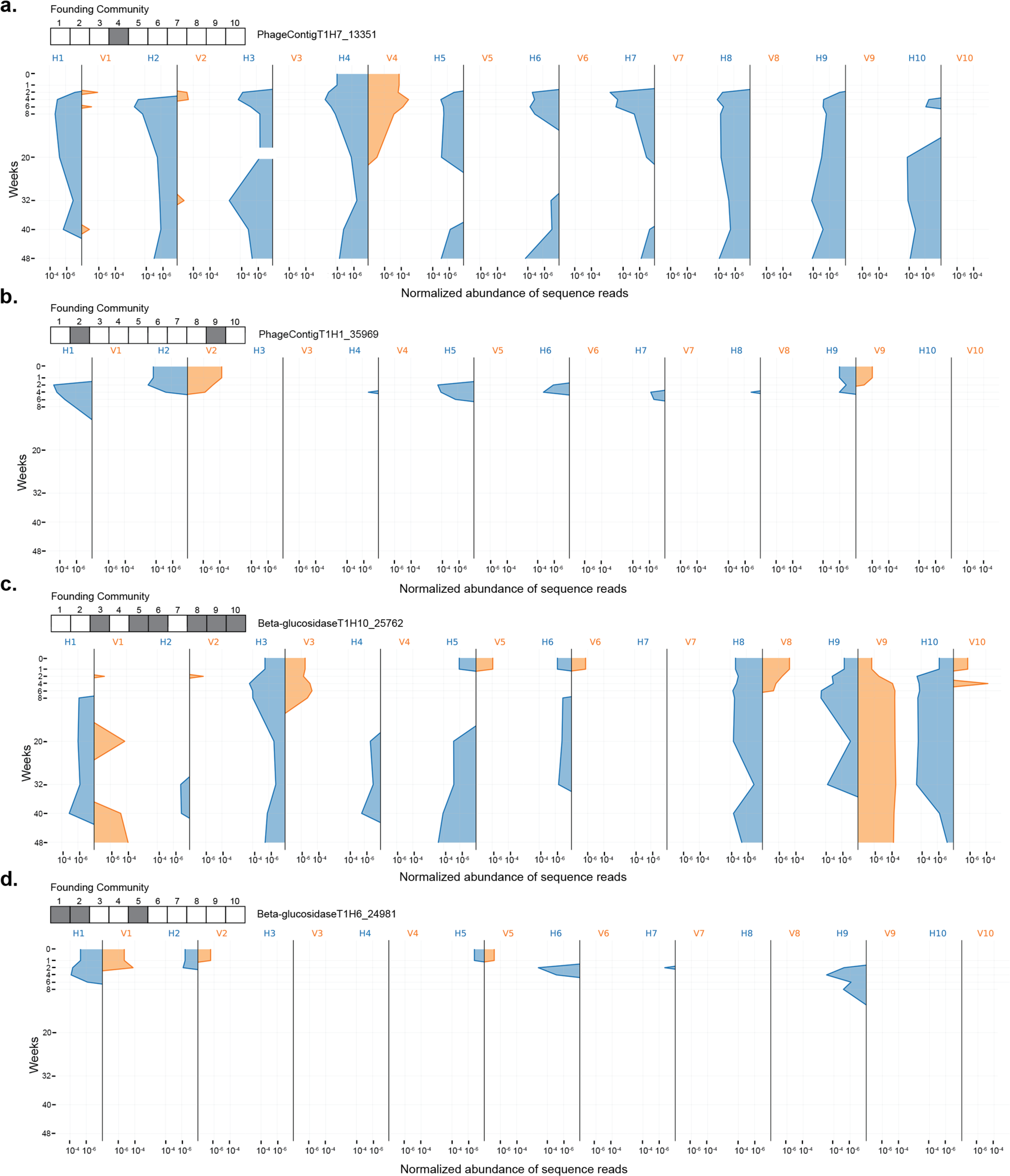
Dynamics of contigs unique to horizontal communities. The dynamics of two-phage like elements (**a**, **b**) and two elements containing predicted beta glucosidases (**c**, **d**) identified from assembling unique reads from horizontal communities at time point T_1_. Each mesocosm is represented as a vertical line and dynamics were tracked through the course of the 48 week experiment. The butterfly plot depicts normalised abundance of sequence reads mapping to each of the representative contigs from horizontal and vertical mesocosms (blue and orange, respectively). Grey / white boxes above each figure depict presence / absence of sequences from each contig in the founding T_0_ community (before the vertical / horizontal split).

Focussing on PhageContigT_1_H_7__13351 (Fig 4a), this phage-like element was detected only in community C4 at T_0_, but was likely present also in communities C1 and C2, although below the limit of detection. By week 3, the phage-like element was present in all horizontal communities. In vertical communities the element persisted in community C4 four for 22 weeks and transiently in C1 and C2, but was present at 48 weeks in all but one horizontal community. Within horizontal communities a pattern observed on four occasions saw the phage-like element go extinct (or fall below the level of detection), but re-emerge and persist at latter time points.

### Ecologically significant genes

To determine whether the horizontal transfer regime affected the abundance and distribution of functional genes, the MG-RAST database was interrogated with sequence reads from all T_0_ and all T_24_ HCs and VCs (Supplementary Table 7). For the ensuing analysis we have avoided standard statistical testing because horizontal microcosms, by virtue of the movement of SGEs and linked genes among communities, cannot be treated as independent entities. Nonetheless, trends are clearly evident from the data displayed in Fig. 5 (and corresponding Supplementary Figs 6a and 6b), which include for each functional category — for each vertical, horizontal and T_0_ community — median value, interquartile range and full data range.

**Figure 5:**
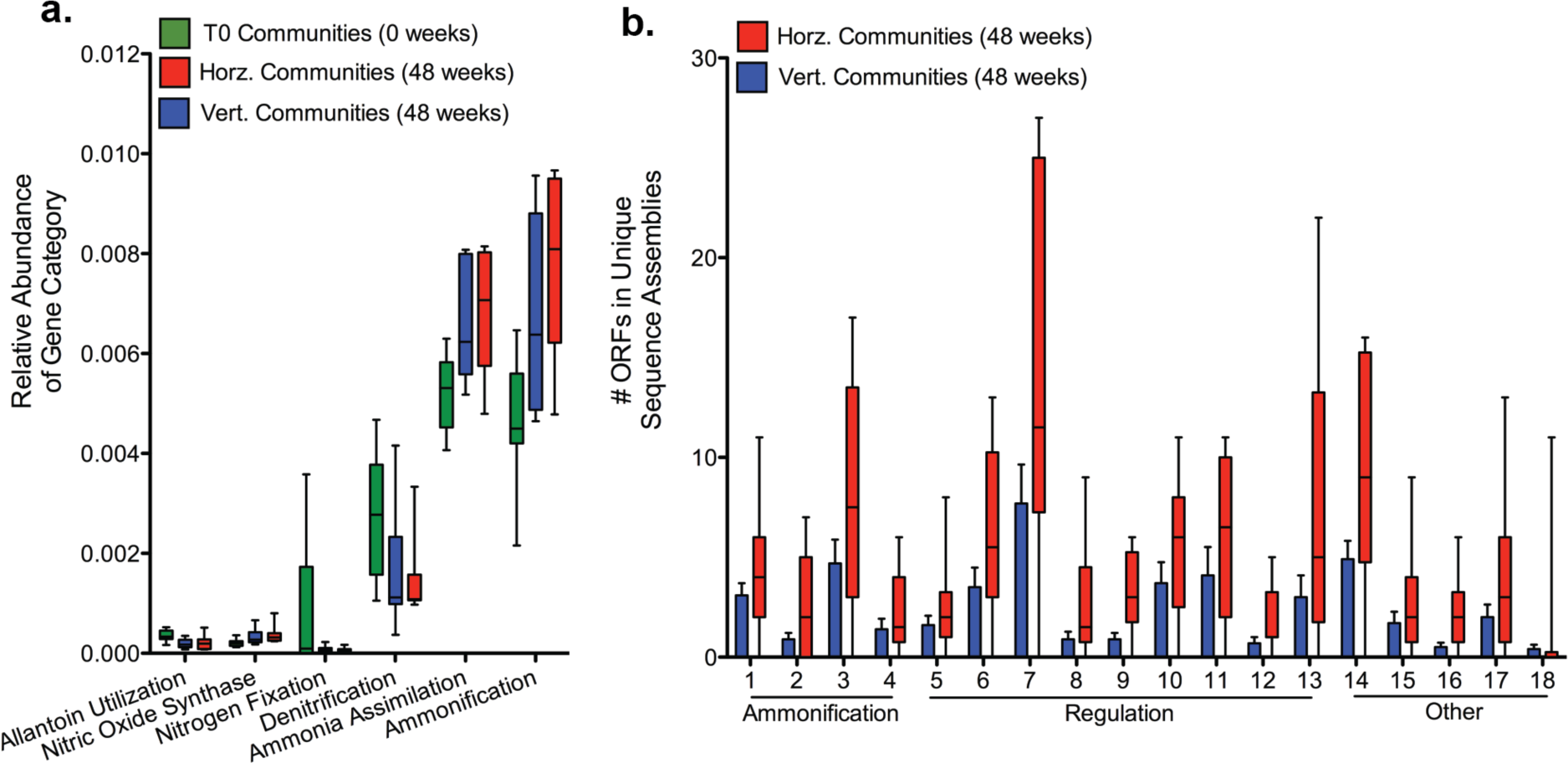
Enrichment of nitrogen metabolism genes in horizontal communities. **a**, Relative abundance of nitrogen metabolism gene categories based on sequence annotation using MG-RAST. Data are shown as box and whisker plots depicting median, inter quartile range (box) and full data spread from 10 replicate communities. **b**, Total number of Open Reading Frames (ORFs) predicted to encode specific domains associated with ammonification, regulation of nitrogen metabolism, and other associated nitrogen metabolism genes in the assemblies of unique vertical and unique horizontal sequences. Data are shown as box and whisker plots depicting median, inter quartile range (box) and full data spread from 10 replicate mesocosms. 1, NifS; 2, NifE; 3, NapA; 4, nitrilase; 5, NtrA; 6, NifR3; 7, NtrY; 8, NifL; 9, PtsN; 10, NtrB; 11, NtrC; 12, NAC; 13, FixI; 14, FixG; 15 & 16 nitroreductases; 17, arginase; 18, cytochrome D1. Full descriptions of domains are provided in Supplementary Table 9.

The relative abundance of reads assigned to 13 of 28 functional categories at Subsystems Level 1 [40], including genes involved in virulence, motility and nitrogen metabolism, changed during the course of the year (Supplementary Fig. 6a). HCs and VCs did not always change to the same extent, or in the same direction (Supplementary Figs 6a and 6b). No obvious change occurred in the overall category of carbohydrate metabolism (Supplementary Fig. 6b), but an increase in *β*-glucosidase metabolism — the rate limiting step in cellulose degradation — was detected in both HCs and VCs (Supplementary Fig. 6a).

The nitrogen-limited nature of the culture medium, combined with evidence of selection favouring genes involved in nitrogen metabolism, led to closer focus on this essential resource. The MG-RAST database was again interrogated, but this time at Subsystems Level 2 [40], which provides information on functional categories within the broader category of nitrogen metabolism (Fig. 5a). Changes were observed in the relative abundance of reads mapping to genes involved in both ammonia assimilation and dissimilation (ammonification). Such changes are consistent with the nature of the selective environment. This prompted a query of the extent to which these changes may have been influenced by movement of genes via SGEs.

A modified version of the bioinformatic pipeline developed to identify sequences unique to HCs (Methods) was applied to the T_24_ metagenomes. T_24_ HC metagenomes were compared to their paired T_24_ VC metagenomes as well as to all metagenomes from VCs from earlier time points (Supplementary Fig. 7a, Supplementary Table 8, Supplementary Methods Fig. 1). This showed a greater number of unique sequences in the HCs, but also showed a fraction of unique sequences present in VCs. The latter is to be expected given that T_0_ communities must contain rare sequences that through the course of the selection experiment became common — and is consistent with data on changes in abundance of genera (Fig. 1). To see whether it was possible to reduce this signal, genomic DNA samples from T_0_ were sequenced on the HiSeq platform resulting in additional 320 million reads per community. The effect of increasing depth of sequence made minimal impact on detection of unique sequences (Supplementary Fig. 7b).Nonetheless, the “deep” T_0_ metagenomes were used in all subsequent analyses.

To determine the fraction of unique reads that mapped to genes involved in nitrogen metabolism, the final sets of unique reads were obtained from HCs and VCs; they were assembled, ORFs identified and functionally categorised using the CDD. HCs contained more unique ORFs predicted to be involved in nitrogen metabolism compared to VCs (Supplementary Fig. 8). HCs were enriched in several functional classes of gene, especially those with predicted roles in regulation and ammonification (Fig. 5b, Supplementary Table 9).

To link to function, we asked whether there was evidence of an effect of the horizontal treatment on a measurable community property. To this end, and with focus on the process of ammonification (production of ammonia through either fixation, or reduction of nitrate / nitrite) the concentration of nitrate, nitrite, and ammonia were determined in the T_24_ HCs and VCs at the end of the two-week period immediately prior to serial passage (Fig. 6). Bearing in mind the non-independence of horizontal communities, two different, but appropriate statistical tests were employed. The first, a two-sample Kolmogorov-Smirnov (K-S) test revealed no significant difference among HCs and VCs for nitrate or nitrite production, but a highly significant difference in the amount of ammonia produced. The second, a Linear Mixed Effect Model, with “community” assigned as a random effect and regime (vertical / horizontal) as a fixed effect, produced results that mirrored those of the K-S test. Details of the tests are provided in the caption to Fig. 6.

**Figure 6:**
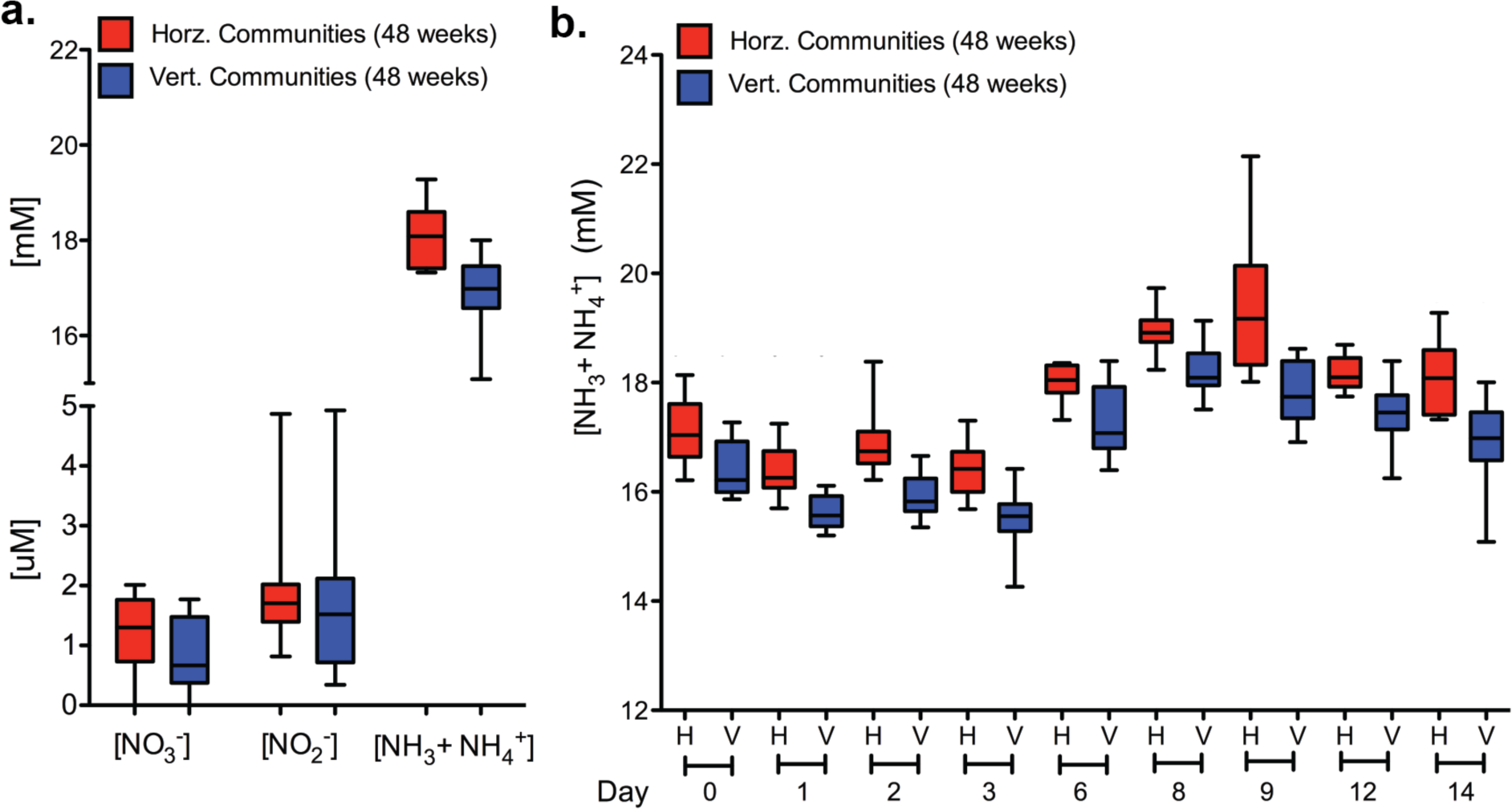
Enhanced ammonia production in horizontal communities. **a**, Concentrations of nitrate, nitrite, and ammonia / ammonium from mesocosms subject to horizontal and vertical regimes at 48 weeks (T_24_). Data are shown as box and whisker plots depicting median, inter quartile range (box) and full data spread from 10 replicate communities. To test for differences in nitrate, nitrite and ammonia / ammonium concentrations at time T_24_ for the ten communities in vertical and horizontal regimes (3 independent measures per community) were compared using a two-sample Kolmogorov-Smirnov test (implemented in python 3.7.2, package scipy 1.3.1). Nitrate and nitrite concentration distributions are not significantly different in the horizontal compared to vertical regime (both *P >* 0.1). However, the distribution of ammonia / ammonium concentrations were significantly different (KS-statistic = 0.6, N = 30 per sample, *P* < 0.001). Analysis was repeated using a Linear Mixed Effect Model. Community identity (1 −10) was assigned as a random effect and regime (horizontal / vertical) as a fixed effect (implemented in python 3.7.2, package stats model 0.10.2, restricted maximum likelihood fit). Results are in agreement with the KS-test. Nitrate and nitrite concentrations were not significantly affected by regime (for both Wald-test results, *P* > 0.3). However, the concentration of ammonia / ammonium was significantly higher in the horizontal regime (mean effect = 1.207, SD = 0.151, 95% CI: [0.912, 1.503], N = 60, Wald-test, *P* < 0.001). **b**, Ammonia / ammonium concentrations measured at nine time points during the two-week incubation period for communities from horizontal and vertical regimes at 48 weeks. Data are shown as box and whisker plots depicting median, inter quartile range (box) and full data spread from 10 replicate communities (triplicate measures were obtained from each community on each occasion). Analysis of time series data using a mixed ANOVA design showed a significant effect of treatment (horizontal vs. vertical; *P* = 1.96 × 10^−5^) and a significant effect of time (*P =* 2.7 × 10^−24^, after Greenhouse & Geisser correction for sphericity). The interaction between treatment regime and time was not significant *P* = 0.69). Comparison between the vertical regime at 48 (T_24_) and community function at the beginning of the experiment (T_0_) showed no difference (see Supplementary Fig. 9).

To check the robustness of this finding, the concentration of ammonia was measured during the course of a two-week period. Despite substantive variability in composition of the independent communities (Supplementary Fig. 2a,b and Supplementary Fig. 3), a significantly greater concentration of ammonia was detected in HCs at each of nine sampling occasions throughout the two week period (Fig. 6b, Supplementary Table 11). A comparison between T0 communities and T24 VCs showed no change (Supplementary Fig. 9) supporting the conclusion that movement of ecologically significant genes via SGEs was responsible for the functional difference between horizontal and vertical communities.

## Discussion

Simplification is a powerful scientific approach. For example, detailed insight into the dynamic of adaptive evolution has come from the field of experimental evolution, where single (often asexual) genotypes are propagated under defined laboratory conditions with variation arising solely by *de novo* mutation and where selection is the primary arbiter of mutation success [24, 25]. This minimal experimental design avoids mixing (migration) between populations, sex and recombination, thus allowing observed evolutionary change to be attributed directly to specific mutations that can be connected causally to changes in fitness.

We sought to apply the same kind of simplifying experimental approach to the study of microbial communities, however doing so is not straightforward. Simply substituting a community of microbes in place of a single bacterial genotype followed by recapitulation of a standard serial transfer experiment, where the community is propagated on a simple carbon source, such as glucose, would yield little insight into processes relevant to natural communities and may even add confusion. In a serial transfer experiment, focal populations adapt genetically by natural selection. Substitution of communities for single populations does not result in communities adapting genetically by natural selection: individual bacterial lineages within the community will evolve — although the extent to which their evolution is adaptive is an open and fascinating question [41–43] — but it makes no sense to think about community-level adaptation, because communities (in a standard serial transfer experiment) are not units of selection. To observe adaptation at the community level, communities must participate in the process of evolution by natural selection. This requires establishment of an experimental regime in which communities are treated as evolving lineages subject to a birth-death process [25, 44–47].

If it makes no sense to assume that communities adapt in a manner analogous to single bacterial populations, then attention shifts away from focus on mutation and selection, and turns to possibilities that might stem from manipulation of evolutionary processes such as recombination, migration [15], or genetic drift. We chose to manipulate the former, seeing possibility to promote or retard HGT via a simple mixing regime predicted to affect the evolutionary fate of SGEs, which in turn, might connect genes to community function.

Before proceeding to discuss choice of experimental design, we note subtleties and complications concerning processes driving eco-evolutionary dynamics, the correspondence between these dynamics and diversity, and the connection between scale of the dynamic and impact on community function. While much is unknown, it seems reasonable to presume a connection, which suggests that communities composed by too few types may fail to capture processes intrinsic to the system, particularly if that process relies on HGT.

Such considerations were instrumental in the decision to found experimental communities from a highly diverse source (garden compost) and to propagate communities on cellulose as sole carbon source. For a start, garden compost is rich in cellulose thus ensuring an *in vitro* environment not too dissimilar from that encountered in nature, additionally, there exists, in principle, the possibility that, in future work, component species might be isolated and cultured on laboratory agar with glucose as sole carbon source (glucose being the utilisable metabolic break-down product of cellulose). However, the primary consideration for use of cellulose was that it is difficult to degrade [27], thus allowing the possibility that communities might be cultured over prolonged periods under nutrient limitation without dominance by *r*-selected types. In effect, while the means and mode of propagation involved serial transfer, communities likely experienced culture conditions more reminiscent of those provided in chemostat culture.

Levels of diversity recorded from analysis of metagenomic data were remarkably high, remaining stable — and even increasing — through the course of the 48 week period of propagation (Fig. 1). This is unexpected given propagation on a single carbon source, where few types should be stably maintained [48–50]. Were the community propagated on glucose instead of cellulose, almost certainly diversity would have been minimal and dominated by a small number of fast growing types. That diversity increased over time (Fig. 1) indicates that rare genera that were initially below the level of detection at the start of the experiment, counter to expectation, increased in frequency. The ecological reasons are not known, but point to unanticipated ecological complexity and presumably abundant opportunity for cross feeding. The high and stably maintained levels of diversity in cellulose-based mesocosms founded from garden compost renders them ideally suited for the kind of experimental work that we have initiated here.

Although noted only in passing, and from examination of the mapping of DNA sequence reads to genes connected with nutrient cycling (data not shown), cycles for replenishing all major nutrients appeared to have been established within each mesocosm, with that involving nitrogen being most evident. Each community harboured a single dominant free-living nitrogen-fixing bacterium, but the identity — even at the genus level — differed between mesocosms. Attempts to establish such complex communities by choosing key players before the fact, would likely fail. That such diversity and stability can arise from a single gram of compost speaks to an abundance of redundancy in these natural systems [51, 52].

Given the experimental design with vertical and horizontal treatment regimes, it is natural to query the relationship between treatment effects and patterns of diversity. At the level of number of bacterial genera and shape of rank abundance curves, no difference was detected and in fact communities showed remarkably similar overall trends. However, differences are apparent when focussing on the rank of individual genera. These are evident in the butterfly plots in Supplementary Fig. 2b and show difference both between independent communities but also differences between vertical and horizontal treatments within communities. Because our experimental design did not include technical replicates (a decision made based on the need to balance experimental feasibility with intuition that we should maximise the number of independent communities) no statistical analysis is possible. Nonetheless, notable are the similarity of changes in genus rankings between vertical and horizontal treatments from the same communities. Also notable is the fact that there was no evidence of convergence in diversity in the horizontal treatment after 48 weeks (Supplementary Fig. 3). Although unsurprising, this emphasises the distinctness of the communities despite being linked via movement of SGEs and linked genes.

The pooling of material from each mesocosm that passed through a 0.2 µm filter to produce a SGE-cocktail, followed by its mixing and regular distribution among HCs, was evidently effective in both mobilising SGEs and promoting horizontal transfer of genes of ecological significance. In fact the magnitude of the effect combined with capacity to detect it via identification of DNA sequence reads unique to each horizontal community, exceeded expectation. Within just two weeks of implementation, ~17 % of DNA sequence reads within HCs were derived from allopatric communities. That these reads could be assembled into contigs containing genes of ecological relevance, including those involved in cellulose degradation, emphasises the dynamism of processes driven by SGEs.

One immediate consequence is utility of the experimental approach to detect — at the level of communities — traits under selection. The suite of unique DNA sequences amplified and disseminated across communities suggests that these genes matter. In the context of conditions experienced by the compost-derived communities this makes sense. Community members are expected to have experienced selection for ability to degrade cellulose: this is indicated by the abundance of horizontally transferred genes predicted to be involved cellulose degradation (Supplementary Fig. 5). Similarly, as mesocosms were nitrogen limited, selection was expected to favour movement of genes involved in nitrogen metabolism. That this happened is indicated by the prevalence of genes implicated in nitrogen metabolism among the set of DNA sequence reads unique to HCs. Not appreciated at the outset was that communities, according to the signature acquired from unique sequence reads, were iron limited. With hindsight this was to be expected. That genes encoding CRISPR are also enriched is especially significant and suggests an unappreciated dynamic between hosts, the defence systems maintained to protect bacterial hosts against phages, and benefits that accrue from the horizontal dissemination of these systems [53]. Application of vertical vs horizontal treatment regimes — which are readily imposed across numerous experimental systems — thus stand to reveal, from the perspective of the community, the nature of traits under selection.

Without further experimentation, the identity of the elements responsible for horizontal spread of genes is unknown, and so too is the relationship between the genes amplified and disseminated across mesocosms, and the vehicles for their dissemination. While some assembled contigs show features typical of phages, many do not. This may be a consequence of working with metagenomic data, but may also reflect the existence of elements whose nature and dynamic remains to be discovered.

One of the larger assembled contigs shown in Supplementary Figure 4b is a case in point: at ~80 kb, hints as to its identity are expected, and yet while carrying genes predicted to encode glycoside hydrolyses — implicated in cellulose degradation — the only suggestion of ability to be both mobilised and disseminated is via linkage to a single integrase. Acquisition of long-read sequence data may aid understanding, but it is also possible that the captured dynamic — the rapid amplification and dissemination of ecologically significant genes — is attributable to “fortunate accidents” mediated by, for example, lytic phages or other elements that happen on rare occasion to capture genes of utility to hosts, with the feedback between genes that prove useful to hosts and the amplifying effects of selection driving their dynamic. This points to the possibility of processes that unfold at the level of diverse communities that remain to be understood.

Irrespective of the nature of the disseminating vehicles, the experimental approach outlined allows the dynamics of elements amplified and disseminated to be tracked. Examples are shown in Figs 3 and 4. Although mechanistic explanations are unavailable, the dynamics of PhageContigT1H7_13351 (Fig. 3A) raise numerous questions. Extinction within VC4, but persistence in all HCs at 48 weeks (with the exception of H1) raises questions as to the causes. Using recently developed Hi-C [16, 19, 54] and population genetic approaches [55] it ought to be possible, in the future, to disentangle ecological from evolutionary effects. Nonetheless, that persistence is evident in HCs, points to a significant role for SGEs.

The association established between genes involved in nitrogen metabolism and enhanced activity of SGEs in HCs is correlative, but is nonetheless linked to horizontally transferred sequences that map to genes with predicted roles in ammonification. The association is further linked to data that demonstrate a significant effect of the experimental treatment on concentrations of ammonia in HCs (Fig. 6), but no effect (compared to T_0_ communities) in VCs (Supplementary Fig. 9). Additionally, increased ammonia in the HCs is consistent with the prediction that horizontal movement of DNA will increase the rate at which community function improves.

The two processes that generate ammonia: nitrogen fixation and nitrate ammonification, both require environments devoid of oxygen (or mostly so in the case of nitrate ammonification) [28, 56]. It is possible that enhanced metabolic activity associated with cellulose degradation causes lower oxygen conditions at the paper surface (where community members grow as biofilms) compared to the VCs, leading to enhanced production of ammonia. Intriguingly, the cattle industry promotes addition of ammonia to feed because of beneficial effects on digestibility of plant matter in the rumen [57]. This raises an alternate possibility, which is that elevated ammonia levels are similarly beneficial to digestion of cellulose in the experimental mesocosms, and reflect a more rapid response to selection in the HCs.

The genomic era has done much to resolve controversy surrounding the importance of horizontal gene transfer as a driver of evolutionary change [58–60]. Direct evidence of HGT — many facilitated by SGEs — from one organism to another are common [61, 62], but the possibility that the process assumes far greater evolutionary significance when operating within communities, has, to the best of our knowledge, been little considered, let alone directly studied. That the vast diversity of DNA sequence encompassed within complex microbial communities might exist in fluid association with SGEs generating permutations of effects significantly beyond those observed through study of individual SGEs and their hosts, is both interesting and plausible.

Microbial communities are dynamical systems defined by complex ecological interactions and evolutionary feedbacks [1]. The challenges associated with understanding this complexity are immense. Here we have demonstrated the utility of an approach in which we have employed a simple experimental manipulation predicted (and shown) to fuel the evolutionary life of SGEs and thus the process of HGT. In effect, mixing (or not) SGEs among communities promotes (or limits) what might be considered community-level sex. The manipulation affects the abundance and distribution of genes of ecological significance, whose individual dynamics can be followed. Further, the effects are sufficient to connect — at a correlative level — genes to community function. That this involves nothing more than frequent exposure of SGEs to new hosts, combined with metagenomic analysis, means that the experimental strategy can be readily transferred to a wide range of microbial communities, from experimental communities as here, through microbiomes of plants and animals, to naturally occurring marine and terrestrial communities.

## Methods

### Establishment of cellulose-degrading microbial communities

Microbial communities were initially established by placing ten independent 1 g samples of fresh compost into 20 mL of M9 minimal media (the nitrogen source was ammonium chloride (0.935 mM)) supplemented with a 4 cm^2^ piece of cellulose paper (Whatman cellulose filter paper). All incubations were performed in 140 mL sterile glass bottles with loosened screw caps allowing for gas exchange between the community and the environment. Incubation was at room temperature without shaking. Compost was sampled from the Square Theodore-Monod compost heap (Paris, France) in February 2016. During the primary incubation period of two weeks the piece of cellulose was suspended by a wire from the screw cap into the media allowing for paper colonization. After two weeks, the community was transferred by moving the wire-suspended cellulose paper into 20 mL of fresh M9 minimal media that also contained a new 4 cm^2^ piece of cellulose paper in suspension. Two more weeks of incubation provided the opportunity for colonization of the new piece of suspended cellulose paper thus eliminating the need for a wire-suspended community for each transfer. All community transfers were performed by vortexing each bottle at maximum speed until the cellulose fibres were disaggregated into a slurry.

### Horizontal and vertical transfer regimes

Vertical and horizontal transfers were performed at the exact same time every two weeks. Before each transfer aliquots of the community were taken for glycerol stocks, DNA extraction, and phage storage. Glycerol stocks were created by mixing 500 µl of cellulose-microbial slurry with 500 µl of 80% glycerol and stored at −80 ºC. For DNA extractions 2 mL of cellulose-microbial slurry was centrifuged at 13,000 g for 10 min and the pellet was stored at −80 ºC for later processing. DNA was extracted from the founding T_0_ communities (before the horizontal / vertical split) as well as transfers 1, 2, 3, 4, 10, 16, 20, 24 (T_1_, T_2_, T_3_, T_4_, T_10_, T_16_, T_20_, T_24_) using the soil DNA extraction kit (Norgen Biotek). DNA sequencing was performed using the MiSeq and NextSeq platforms for all time points and T_0_ communities with additional deep sequencing of the T_0_ communities using the HiSeq platform.

Transfers at two-week intervals were performed as described in Fig. 2. The “SGE-cocktail” was prepared by mixing 1 mL of homogenized community (n=10) followed by centrifugation and passage through a 0.2 µm filter to remove microbial cells. Recognising the possibility that ultra micro-bacteria might escape sedimentation and pass through the filter, metagenomic sequence reads were recruited to seven available reference genomes. A maximum of 0.018 % of reads across all horizontal and vertical mesocosms at 48 weeks matched these reference genomes (Supplementary Fig. 10).

### Bioinformatic analysis

DNA sequences were demultiplexed using bcl2fastq, paired ends were joined using FLASh [63] and initial fastq files were generated. Preprocessing of fastq files was performed using PrinSeq [64] with a minimum length of 100 base pairs, minimum quality score of 25, and a maximum percentage of N’s of 10%. All metagenomes were uploaded to the MG-RAST metagenomic analysis server and are publicly available (Supplementary Table 1). Gene category abundances (Supplementary Fig. 5) and genus identifications (Supplementary Table 6) were determined using the MG-RAST pipeline [65]. Sequence matches were determined using BLASTn [66] with a minimum e-value threshold of 1E-05, minimum alignment length of 100 base pairs, and a minimum percentage identity of 90%. “Unique” Horizontal and Vertical sequences were assembled using MEGAHIT [67], ORFs with a minimum length of 100 amino acids were extracted using getORF [68], compared to the Conserved Domain Database (CDD) [35] using RPS-BLAST, and top domain hits were extracted (Supplementary Table 4). Phage-like contigs were classified based on phage-specific protein domains (Supplementary Table 3) and their distribution was determined using Fr-Hit [69]. Sequences matches to phage-like contig were based on a minimum alignment length of 100 base pairs with a percentage identity of at least 90%. Nitrogen-related genes were based on gene function classifications and descriptions with a total of 136 domains (Supplementary Table 9). To identify unique sequences a custom bioinformatic pipeline was developed (Supplementary Methods Fig. 1). Each Horizontal metagenome was compared to the founding T_0_ metagenome (before the Horizontal vs. Vertical split) and the paired Vertical metagenome using BLASTn. Horizontal-T_0_ and Horizontal-Vertical sequence matches were removed, thus leaving a set of unique Horizontal sequences that could not be identified in neither the founding T_0_-metagenome, nor the paired Vertical-metagenome. Sequence matches were determined using a minimum e-value threshold of 1E-05, minimum alignment length of 100 base pairs, and a minimum percent identity of 90%. The unique horizontal sequences were then assembled, Open Reading Frames (ORFs) were extracted, and predicted protein domains were determined through comparison to the Conserved Domain Database (CDD). To determine the impact of increasing the sequencing depth of the T_0_ communities on the final percentage of unique sequences, T_0_ metagenomes were sequenced using the HiSeq platform and reads from each mesocosm split arbitrarily into eight metagenomes each consisting of about ~40 million sequences. The unique analysis pipeline was then applied using the first arbitrary T_0_ metagenome and any matching sequences were removed. The process was repeated with all eight T_0_ arbitrarily defined metagenomes thus revealing the extent to which additional sequencing affected ability to detect rare sequences.

### Nitrogen assays

Vertical and Horizontal communities from the one-year time point (24 transfers, T_24_) and T_0_ were re-established by placing 100 µl of glycerol stock into 20 mL of M9 minimal media supplemented with a 4 cm^2^ piece of cellulose paper and incubated for two weeks. 1 mL of the cellulose-community slurry was transferred to 19 mL of M9 Minimal Media with a fresh piece of cellulose and incubated for an additional two weeks. 1 mL of the community was then transferred to a new bottle and 100 µl of surrounding media was sampled at various time points during the two-week incubation period to determine ammonia/ammonium, nitrate, and nitrite concentrations. The concentrations of all three nitrogen species were determined using fluorometric assay kits according the manufacturers protocol (Ammonia Assay Kit, Sigma-Alrdrich, and nitrate/nitrite fluorometric assay kit) (Fig. 6 and Supplementary Table 10).

## Data availability

All metagenomes are publicly available here: https://www.mg-rast.org/mgmain.html?mgpage=project&project=mgp18485. Supplementary Table 1 contains metagenome descriptions and MG-RAST identifiers

All associated datasets are publicly available here: https://zenodo.org/record/3293312 and Supplementary Table 12 contains file descriptions and associated metadata

## Supplementary files

Supplementary Figure 1: Changes in diversity.

Supplementary Figure 2: Changes in patterns of diversity seen in rank abundance plots.

Supplementary Figure 3: Among community diversity.

Supplementary Figure 4: Contigs assembled from unique sequence reads.

Supplementary FFigure 5: Ecologically significant genes.

Supplementary Figure 6: Functional gene categories.

Supplementary Figure 7: Distinguishing sequences transferred by SGEs from rare sequences.

Supplementary Figure 8: Nitrogen-related genes are enriched in “unique” horizontal sequences.

Supplementary Figure 9: No change in function of communities subject to the vertical transfer regime after 48 weeks.

Supplementary Figure 10: DNA sequence reads mapping to ultra-microbacteria reference genomes.

Supplementary Methods Figure 1: Bioinformatic pipeline for identifying “unique” sequences.

Supplementary Table 1: Genus abundance of all sequenced time points for all mesocosms based on the MG-RAST annotation pipeline.

Supplementary Table 2: MGRAST IDs and metadata associated with metagenomes.

Supplementary Table 3: Summary of unique sequences assembled from T_1_ Horizontal mesocosms.

Supplementary Table 4: Sequences matching to phage-like contigs in Fig. 2; annotation of phage-like contigs depicted in Fig. 2; all T_1_ unique contigs containing an ORF associated with bacteriophages.

Supplementary Table 5: Number of unique T_1_ horizontal sequences containing an ORF with a domain of interest depicted in Supplementary Fig. 2.

Supplementary Table 6: Full annotation of the two phage-like and two beta-glucosidase contigs depicted in Fig. 4.

Supplementary Table 7: The relative abundance of functional gene categories in T_0_, T_24_ vertical, and T_24_ horizontal metagenomes based on the MG-RAST annotation pipeline.

Supplementary Table 8: Summary of unique sequences assembled from T_24_ horizontal and T_24_ vertical metagenomes.

Supplementary Table 9: Number of nitrogen-related ORFs assembled from unique T_24_ horizontal sequences depicted in Fig. 5b; number of nitrogen-related ORFs assembled from unique T24 horizontal sequences.

Supplementary Table 10: Nitrate, nitrite, and ammonia / ammonium concentrations of T_24_ horizontal and T24 vertical mesocosms at the end of the two-week incubation period depicted in Figure 6a.

Supplementary Table 11: Time series of ammonia / ammonium concentrations of T_24_ horizontal and T_24_ vertical mesocosms during the two-week incubation period.

Supplementary Table 12: Description of all associated datasets publicly available at https://zenodo.org/record/3293312.

## Acknowledgements

We thank members of the Rainey lab at ESPCI and the MPI for Evolutionary Biology, and especially Dave Rogers and Andy Farr, for valuable discussion, Boris Shraiman and Daniel Fisher for fuelling interest in the intractability of microbial communities, Sven Kuenzel for DNA sequencing, and Rob Edwards and Jeremy Barr for comments on the manuscript. S.Q. acknowledges receipt of funding from the European Union’s Horizon 2020 research and innovation programme under the Marie Skłodowska-Curie grant agreement No 747527.

## Author information

Concept & experimental design PBR & SQ, experimentation SQ & PBR, bioinformatic analysis SQ and GD, writing PBR & SQ.

## References

[1] Widder, S., Allen, R. J., Pfeiffer, T., Curtis, T. P., Wiuf, C., Sloan, W. T., Cordero, O. X., Brown, S. P., Momeni, B., Shou, W., et al. 2016 Challenges in microbial ecology: building predictive understanding of community function and dynamics. ISME J. 10, 2557–2568. (DOI: 10.1038/ismej.2016.45).

[2] Knight, R., Callewaert, C., Marotz, C., Hyde, E. R., Debelius, J. W., McDonald, D. & Sogin, M. L. 2017 The microbiome and human biology. Annu Rev Genomics Hum Genet 18, 65–86. (DOI: 10.1146/annurev-genom-083115-022438).

[3] Gilbert, J. A., Blaser, M. J., Caporaso, J. G., Jansson, J. K., Lynch, S. V. & Knight, R. 2018 Current understanding of the human microbiome. Nat Med 24, 392–400. (DOI: 10.1038/nm.4517).

[4] Tyson, G. W., Chapman, J., Hugenholtz, P., Allen, E. E., Ram, R. J., Richardson, P. M., Solovyev, V. V., Rubin, E. M., Rokhsar, D. S. & Banfield, J. F. 2004 Community structure and metabolism through reconstruction of microbial genomes from the environment. Nature 428, 37–43. (DOI: 10.1038/nature02340).

[5] Allison, S. D. & Martiny, J. B. 2008 Colloquium paper: resistance, resilience, and redundancy in microbial communities. Proc Natl Acad Sci U S A 105 Suppl 1, 11512–11519. (DOI: 10.1073/pnas.0801925105).

[6] Fuhrman, J. A., Cram, J. A. & Needham, D. M. 2015 Marine microbial community dynamics and their ecological interpretation. Nat Rev Microbiol 13, 133–146. (DOI: 10.1038/nrmicro3417).

[7] Anantharaman, K., Brown, C. T., Hug, L. A., Sharon, I., Castelle, C. J., Probst, A. J., Thomas, B. C., Singh, A., Wilkins, M. J., Karaoz, U., et al. 2016 Thousands of microbial genomes shed light on interconnected biogeochemical processes in an aquifer system. Nat Commun 7, 13219. (DOI: ARTN 13219 10.1038/ncomms13219).

[8] Bardgett, R. D., Freeman, C. & Ostle, N. J. 2008 Microbial contributions to climate change through carbon cycle feedbacks. ISME J 2, 805–814. (DOI: 10.1038/ismej.2008.58).

[9] Venter, J. C., Remington, K., Heidelberg, J. F., Halpern, A. L., Rusch, D., Eisen, J. A., Wu, D. Y., Paulsen, I., Nelson, K. E., Nelson, W., et al. 2004 Environmental genome shotgun sequencing of the Sargasso Sea. Science 304, 66–74. (DOI: 10.1126/science.1093857).

[10] Quince, C., Walker, A. W., Simpson, J. T., Loman, N. J. & Segata, N. 2017 Shotgun metagenomics, from sampling to analysis. Nat Biotechnol 35, 833–844. (DOI: 10.1038/nbt.3935).

[11] Koskella, B., Hall, L. J. & Metcalf, C. J. E. 2017 The microbiome beyond the horizon of ecological and evolutionary theory. Nat Ecol Evol 1, 1606–1615. (DOI: 10.1038/s41559-017-0340-2).

[12] Hansen, S. K., Rainey, P. B., Haagensen, J. A. & Molin, S. 2007 Evolution of species interactions in a biofilm community. Nature 445, 533–536.

[13] Lawrence, D., Fiegna, F., Behrends, V., Bundy, J. G., Phillimore, A. B., Bell, T. & Barraclough, T. G. 2012 Species interactions alter evolutionary responses to a novel environment. PLoS Biol 10, e1001330. (DOI: 10.1371/journal.pbio.1001330).

[14] Hug, L. A. & Co, R. 2018 It takes a village: Microbial communities thrive through interactions and metabolic handoffs. mSystems 3, e00152–00117. (DOI: ARTN e00152-17 10.1128/mSystems.00152-17).

[15] Robinson, C. D., Klein, H. S., Murphy, K. D., Parthasarathy, R., Guillemin, K. & Bohannan, B. J. M. 2018 Experimental bacterial adaptation to the zebrafish gut reveals a primary role for immigration. PLoS Biol. 16. (DOI: ARTN e2006893 10.1371/journal.pbio.2006893).

[16] Marbouty, M., Baudry, L., Cournac, A. & Koszul, R. 2017 Scaffolding bacterial genomes and probing host-virus interactions in gut microbiome by proximity ligation (chromosome capture) assay. Sci. Adv. 3, e1602105. (DOI: 10.1126/sciadv.1602105).

[17] Truong, D. T., Tett, A., Pasolli, E., Huttenhower, C. & Segata, N. 2017 Microbial strain-level population structure and genetic diversity from metagenomes. Genome Res 27, 626–638. (DOI: 10.1101/gr.216242.116).

[18] Garud, N. R., Good, B. H., Hallatschek, O. & Pollard, K. S. 2019 Evolutionary dynamics of bacteria in the gut microbiome within and across hosts. PLoS Biol 17, e3000102. (DOI: 10.1371/journal.pbio.3000102).

[19] Yaffe, E. & Relman, D. A. 2019 Tracking microbial evolution in the human gut using Hi-C. BioRxiv, http://dx.doi.org/10.1101/594903.

[20] Doolittle, W. F. & Sapienza, C. 1980 Selfish genes, the phenotype paradigm and genome evolution. Nature 284, 601–603.

[21] Orgel, L. E. & Crick, F. H. C. 1980 Selfish DNA: The ultimate parasite. Nature 284, 604–607.

[22] Bergstrom, C. T., Lipsitch, M. & Levin, B. R. 2000 Natural selection, infectious transfer and the existence conditions for bacterial plasmids. Genetics 155, 1505–1519.

[23] Burt, A. & Trivers, R. L. 2006 Genes in Conflict: the Biology of Selfish Genetic Elements. Harvard, Cambridge, MA., Belknap Press.

[24] Elena, S. F. & Lenski, R. E. 2003 Evolution experiments with microorganisms: the dynamics and genetic bases of adaptation. Nat Rev Genet 4, 457–469.

[25] Rainey, P. B., Remigi, P., Farr, A. D. & Lind, P. A. 2017 Darwin was right: where now for experimental evolution. Curr Opin Genet Dev 47, 102–109. (DOI: 10.1016/j.gde.2017.09.003).

[26] Maltez Thomas, A., Prata Lima, F., Maria Silva Moura, L., Maria da Silva, A., Dias-Neto, E. & Setubal, J. C. 2018 Comparative Metagenomics. Methods Mol Biol 1704, 243–260. (DOI: 10.1007/978-1-4939-7463-4_8).

[27] Wilson, D. B. 2011 Microbial diversity of cellulose hydrolysis. Curr Opin Microbiol 14, 259–263. (DOI: 10.1016/j.mib.2011.04.004).

[28] Simon, J. 2002 Enzymology and bioenergetics of respiratory nitrite ammonification. FEMS Microbiol Rev 26, 285–309. (DOI: 10.1111/j.1574-6976.2002.tb00616.x).

[29] Goddard, M. R., Godfray, H. C. J. & Burt, A. 2005 Sex increases the efficacy of natural selection in experimental yeast populations. Nature 434, 636–640. (DOI: 10.1038/nature03405).

[30] McDonald, M. J., Rice, D. P. & Desai, M. M. 2016 Sex speeds adaptation by altering the dynamics of molecular evolution. Nature 531, 233–+. (DOI: 10.1038/nature17143).

[31] Aminov, R. I. 2011 Horizontal gene exchange in environmental microbiota. Front Microbiol 2, 158. (DOI: 10.3389/fmicb.2011.00158).

[32] Colombi, E., Straub, C., Kunzel, S., Templeton, M. D., McCann, H. C. & Rainey, P. B. 2017 Evolution of copper resistance in the kiwifruit pathogen Pseudomonas syringae pv. actinidiae through acquisition of integrative conjugative elements and plasmids. Environ Microbiol 19, 819–832. (DOI: 10.1111/1462-2920.13662).

[33] Hall, J. P. J., Brockhurst, M. A. & Harrison, E. 2017 Sampling the mobile gene pool: innovation via horizontal gene transfer in bacteria. Phil Trans R Soc Lond B 372. (DOI: 10.1098/rstb.2016.0424).

[34] Casjens, S. 2003 Prophages and bacterial genomics: what have we learned so far? Mol. Microbiol. 49, 277–300. (DOI: 10.1046/j.1365-2958.2003.03580.x).

[35] Marchler-Bauer, A., Derbyshire, M. K., Gonzales, N. R., Lu, S., Chitsaz, F., Geer, L. Y., Geer, R. C., He, J., Gwadz, M., Hurwitz, D. I., et al. 2015 CDD: NCBI’s conserved domain database. Nucleic Acids Res. 43, D222–226. (DOI: 10.1093/nar/gku1221).

[36] Seed, K. D., Lazinski, D. W., Calderwood, S. B. & Camilli, A. 2013 A bacteriophage encodes its own CRISPR/Cas adaptive response to evade host innate immunity. Nature 494, 489–491. (DOI: 10.1038/nature11927).

[37] Ramisetty, B. C. & Santhosh, R. S. 2016 Horizontal gene transfer of chromosomal Type II toxin-antitoxin systems of Escherichia coli. FEMS Microbiol Lett 363. (DOI: 10.1093/femsle/fnv238).

[38] Bustamante, P. & Iredell, J. R. 2017 Carriage of type II toxin-antitoxin systems by the growing group of IncX plasmids. Plasmid 91, 19–27. (DOI: 10.1016/j.plasmid.2017.02.006).

[39] Singhania, R. R., Patel, A. K., Sukumaran, R. K., Larroche, C. & Pandey, A. 2013 Role and significance of beta-glucosidases in the hydrolysis of cellulose for bioethanol production. Bioresour Technol 127, 500–507. (DOI: 10.1016/j.biortech.2012.09.012).

[40] Overbeek, R., Begley, T., Butler, R. M., Choudhuri, J. V., Chuang, H. Y., Cohoon, M., de Crecy-Lagard, V., Diaz, N., Disz, T., Edwards, R., et al. 2005 The subsystems approach to genome annotation and its use in the project to annotate 1000 genomes. Nucleic Acids Res. 33, 5691–5702. (DOI: 10.1093/nar/gki866).

[41] Chu, H. Y., Sprouffske, K. & Wagner, A. 2018 Assessing the benefits of horizontal gene transfer by laboratory evolution and genome sequencing. BMC Evol. Biol. 18, 54. (DOI: 10.1186/s12862-018-1164-7).

[42] Frazão, N., Sousa, A., Lässig, M. & Gordo, I. 2019 Horizontal gene transfer overrides mutation in Escherichia coli colonizing the mammalian gut. Proc Natl Acad Sci USA, 201906958. (DOI: 10.1073/pnas.1906958116).

[43] Zhao, S. J., Lieberman, T. D., Poyet, M., Kauffman, K. M., Gibbons, S. M., Groussin, M., Xavier, R. J. & Alm, E. J. 2019 Adaptive evolution within gut microbiomes of healthy people. Cell Host & Microbe 25, 656–667. (DOI: 10.1016/j.chom.2019.03.007).

[44] Wilson, D. S. & Sober, E. 1989 Reviving the superorganism. J Theor Bio. 136, 337–356.

[45] Swenson, W., Wilson, D. S. & Elias, R. 2000 Artificial ecosystem selection. Proc Natl Acad Sci U S A 97, 9110–9114. (DOI: 10.1073/pnas.150237597 150237597).

[46] Xie, L., Yuan, A. E. & Shou, W. Y. 2019 Simulations reveal challenges to artificial community selection and possible strategies for success. PLoS Biol. 17. (DOI: ARTN e3000295 10.1371/journal.pbio.3000295).

[47] Black, A. J., Bourrat, P. & Rainey, P. B. 2020 Ecological scaffolding and the evolution of individuality. Nat Ecol Evol In press.

[48] Gause, G. F. 1934 The Struggle for Existence. Baltimore, Williams & Wilkins.

[49] Rosenzweig, R. F., Sharp, R. R., Treves, D. S. & Adams, J. 1994 Microbial evolution in a simple unstructured environment: Genetic differentiation in *Escherichia coli*. Genetics 137, 903–917.

[50] Rainey, P. B., Buckling, A., Kassen, R. & Travisano, M. 2000 The emergence and maintenance of diversity: insights from experimental bacterial populations. Trends Ecol Evol 15, 243–247.

[51] Rosenfeld, J. S. 2002 Functional redundancy in ecology and conservation. Oikos 98, 156–162.

[52] Louca, S., Polz, M. F., Mazel, F., Albright, M. B. N., Huber, J. A., O’Connor, M. I., Ackermann, M., Hahn, A. S., Srivastava, D. S., Crowe, S. A., et al. 2018 Function and functional redundancy in microbial systems. Nat Ecol Evol 2, 936–943. (DOI: 10.1038/s41559-018-0519-1).

[53] Landsberger, M., Gandon, S., Meaden, S., Rollie, C., Chevallereau, A., Buckling, A., Westra, E. R. & van Houte, S. 2018 Anti-CRISPR phages cooperate to overcome CRISPR-Cas immunity. Cell 174, 908–916. (DOI: 10.1016/j.cell.2018.05.058).

[54] Marbouty, M., Cournac, A., Flot, J. F., Marie-Nelly, H., Mozziconacci, J. & Koszul, R. 2014 Metagenomic chromosome conformation capture (meta3C) unveils the diversity of chromosome organization in microorganisms. eLife 3, e03318. (DOI: 10.7554/eLife.03318).

[55] Garud, N. R., Good, B. H., Hallatschek, O. & Pollard, K. S. 2019 Evolutionary dynamics of bacteria in the gut microbiome within and across hosts. PLoS Biol. 17, e3000102. (DOI: 10.1371/journal.pbio.3000102).

[56] Lam, P. & Kuypers, M. M. 2011 Microbial nitrogen cycling processes in oxygen minimum zones. Ann Rev Mar Sci 3, 317–345. (DOI: 10.1146/annurev-marine-120709-142814).

[57] Givens, D. I., Adamson, A. H. & Cobby, J. M. 1988 The effect of ammoniation on the nutritive value of wheat, barley and oat straws. II. Digestibility and energy value measurements in vivo and their prediction from laboratory measurements. Anim Feed Sci Tech 19, 173–184.

[58] Goldenfeld, N. & Woese, C. 2007 Biology’s next revolution. Nature 445, 369. (DOI: 10.1038/445369a).

[59] Koonin, E. V. 2009 Darwinian evolution in the light of genomics. Nucleic Acids Res 37, 1011–1034. (DOI: 10.1093/nar/gkp089).

[60] Boto, L. 2010 Horizontal gene transfer in evolution: facts and challenges. Proc Biol Sci 277, 819–827. (DOI: 10.1098/rspb.2009.1679).

[61] Ochman, H., Lawrence, J. G. & Groisman, E. A. 2000 Lateral gene transfer and the nature of bacterial innovation. Nature 405, 299–304.

[62] Werren, J. H. 2011 Selfish genetic elements, genetic conflict, and evolutionary innovation. Proc Natl Acad Sci U S A 108 Suppl 2, 10863–10870. (DOI: 10.1073/pnas.1102343108).

[63] Magoc, T. & Salzberg, S. L. 2011 FLASH: fast length adjustment of short reads to improve genome assemblies. Bioinformatics 27, 2957–2963. (DOI: 10.1093/bioinformatics/btr507).

[64] Schmieder, R. & Edwards, R. 2011 Quality control and preprocessing of metagenomic datasets. Bioinformatics 27, 863–864. (DOI: 10.1093/bioinformatics/btr026).

[65] Wilke, A., Bischof, J., Gerlach, W., Glass, E., Harrison, T., Keegan, K. P., Paczian, T., Trimble, W. L., Bagchi, S., Grama, A., et al. 2016 The MG-RAST metagenomics database and portal in 2015. Nucleic Acids Res. 44, D590–594. (DOI: 10.1093/nar/gkv1322).

[66] Altschul, S. F., Gish, W., Miller, W., Myers, E. W. & Lipman, D. J. 1990 Basic local alignment search tool. J Mol Biol 215, 403–410. (DOI: 10.1016/S0022-2836(05)80360-2).

[67] Li, D., Liu, C. M., Luo, R., Sadakane, K. & Lam, T. W. 2015 MEGAHIT: an ultra-fast single-node solution for large and complex metagenomics assembly via succinct de Bruijn graph. Bioinformatics 31, 1674–1676. (DOI: 10.1093/bioinformatics/btv033).

[68] Rice, P., Longden, I. & Bleasby, A. 2000 EMBOSS: the European Molecular Biology Open Software Suite. Trends Genet. 16, 276–277.

[69] Niu, B., Zhu, Z., Fu, L., Wu, S. & Li, W. 2011 FR-HIT, a very fast program to recruit metagenomic reads to homologous reference genomes. Bioinformatics 27, 1704–1705. (DOI: 10.1093/bioinformatics/btr252).

