## Supplementary Information for "Experimental manipulation of selfish genetic elements links genes to microbial community function"

### Supplementary Figures

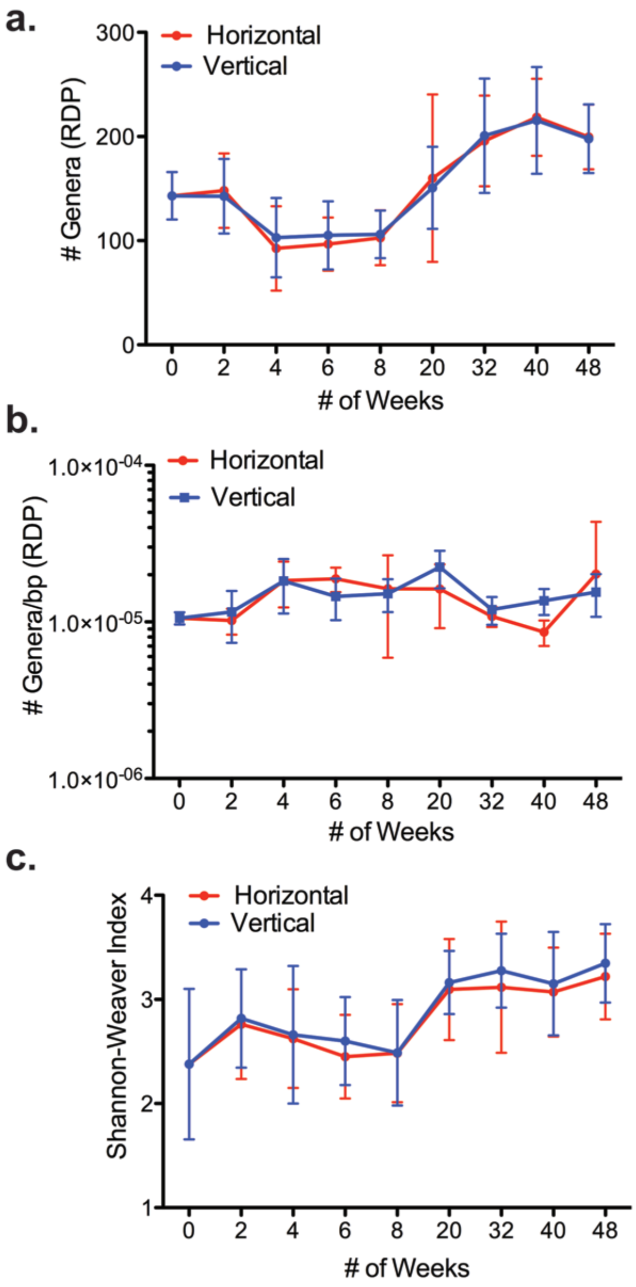


**Supplementary Figure 1: Changes in diversity.**

**a,** The total number of genera based on the 16s rRNA subunit extracted from sequences annotated using the MG-RAST database. **b,** The number of genera normalized to the total number of sequences per metagenome. **c,** Shannon-Weaver diversity index. For all panels data are means and standard deviation of ten independent replicates.

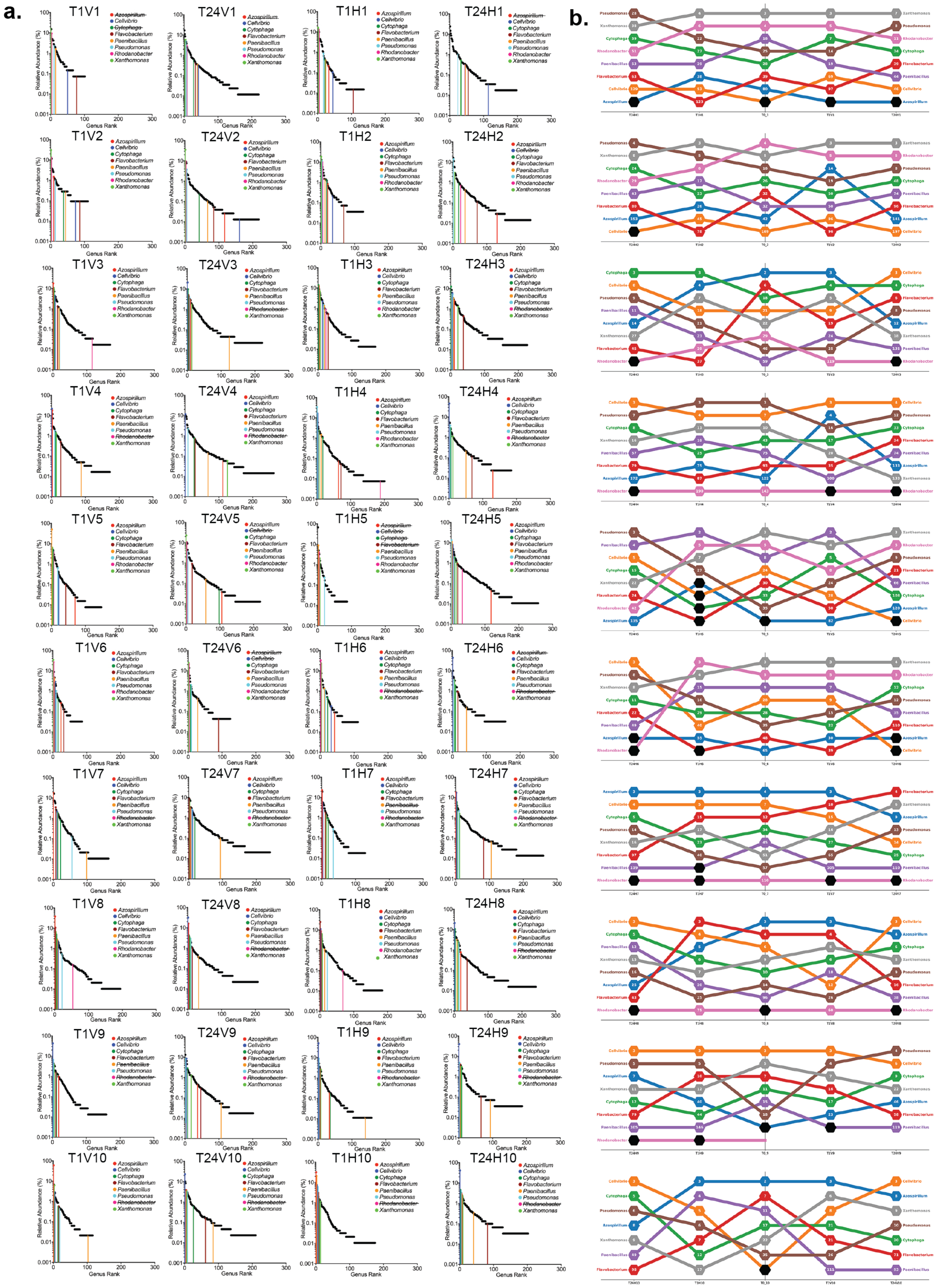
a.
b.
T1V1
T24V1
T1H1
T24H1
T1V2
T24V2
T1H2
T24H2
T1V3
T24V3
T1H3
T24H3
T1V4
T24V4
T1H4
T24H4
T1V5
T24V5
T1H5
T24H5
T1V6
T24V6
T1H6
T24H6
T1V7
T24V7
T1H7
T24H7
T1V8
T24V8
T1H8
T24H8
T1V9
T24V9
T1H9
T24H9
T1V10
T24V10
T1H10
T24H10
Azospirillum
Cellvibrio
Cytophaga
Flavobacterium
Paenibacillus
Pseudomonas
Rhodanobacter
Xanthomonas
Relative Abundance (%)
Genus Rank

**Supplementary Figure 2: Changes in patterns of diversity seen in rank abundance plots**

**a,** Genus-level rank abundance curves for each vertical and horizontal community at time points $T_1$ (2 weeks after the $T_0$ split) and $T_{24}$ (48 weeks). Vertical coloured lines correspond to the rank of the eight most commonly found genera at $T_0$. **b,** Butterfly plots showing changes in the rank abundance of the eight common genera starting with $T_0$ (centre), then $T_1$ horizontal and $V_1$ vertical communities (immediately left and right of centre, respectively) and $T_{24}$ horizontal and $V_{24}$ vertical communities (outer left and right columns, respectively). The numbers in the centre of each node indicate the rank within the respective mesocosm.

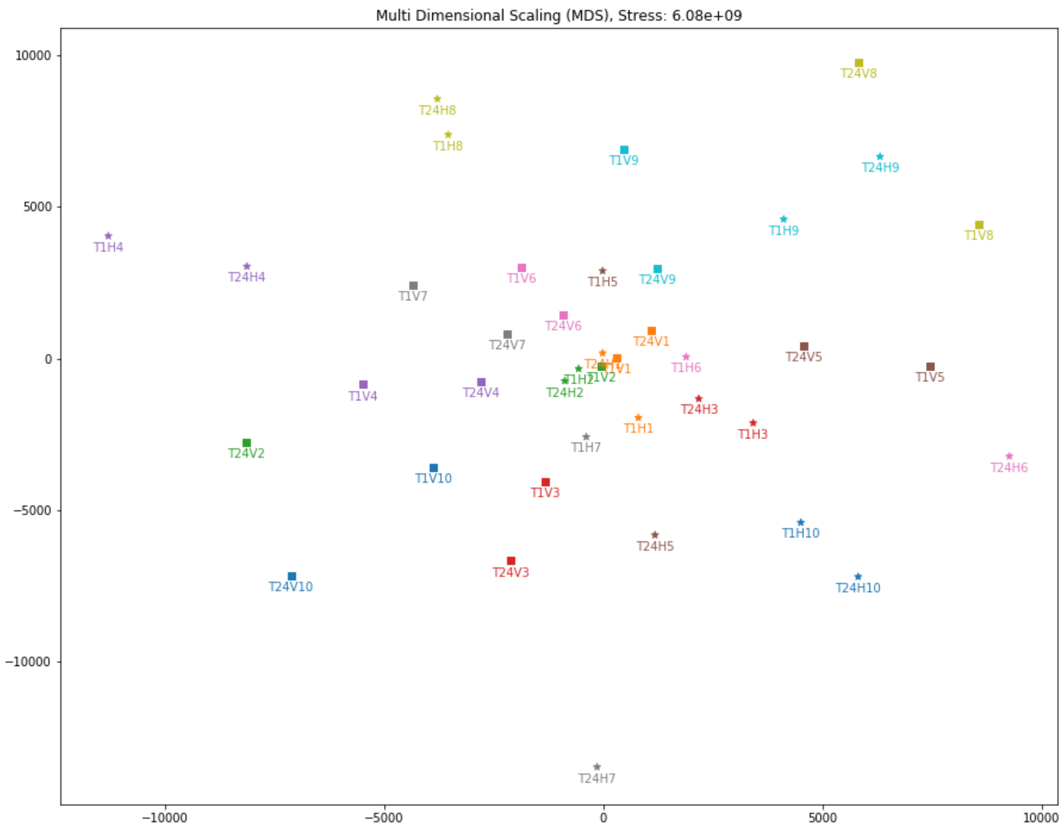


### Supplementary Figure 3: Among community diversity

Multidimensional scaling was used to display differences among communities from time point 1 (T1), two weeks after communities were split into vertical and horizontal treatments, and time point 24 (T24) 48 weeks after the split. Input data are from the 16s rRNA subunit extracted from sequences annotated using the MG-RAST database and categorised at genus level. Communities are distinguished by colour with squares depicting vertical and stars depicting horizontal communities, respectively.

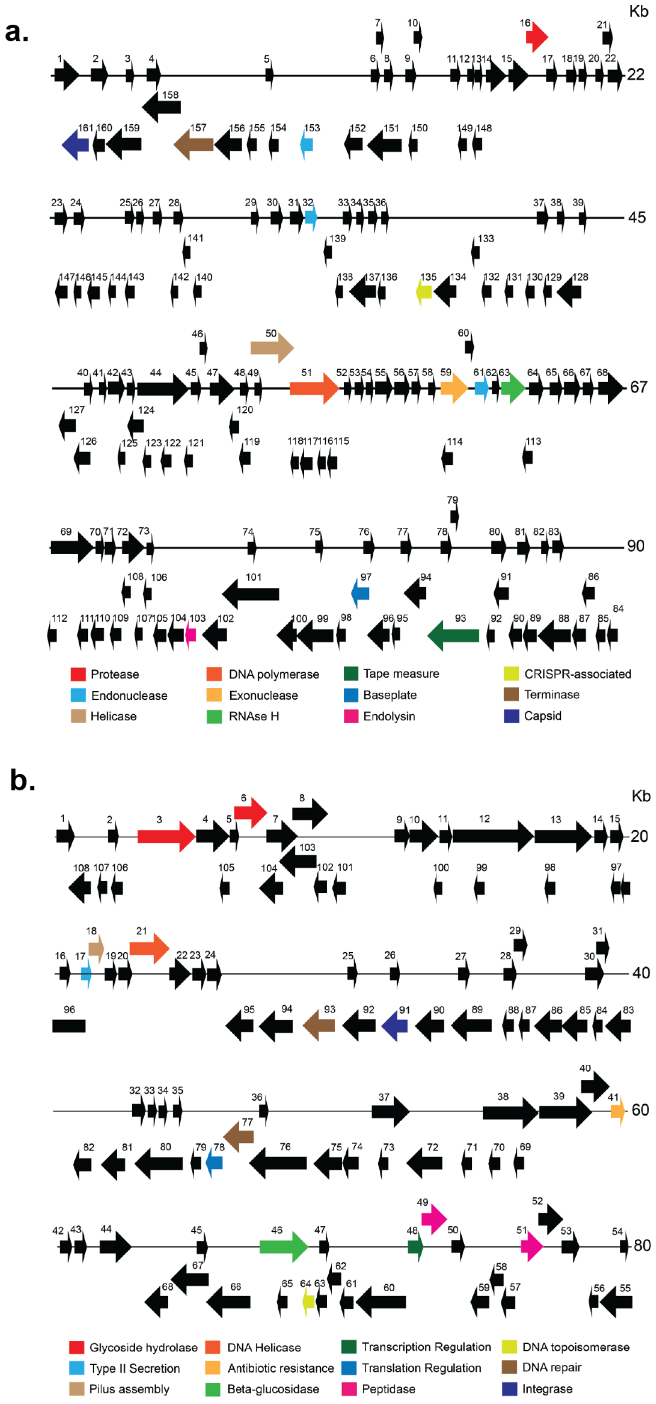


**Supplementary Figure 4: contigs assembled from unique sequence reads.**

Two of more than 3,000 contigs greater than 1 kb assembled from DNA sequence reads uniquely found in horizontal communities at time point $T_1$ and absent from the paired vertical community and from the parental ($T_0$) community. **a,** A ~90 kb contig from $T_1$H-1 containing many phage-specific domains including open-reading frames predicted to encode proteins involved with capsid structure, assembly of phage particles, and bacterial cell lysis. **b,** An ~80 kb contig from $T_1$H-7 containing a predicted cellulose-degrading domain as well as various domains associated with mobile elements. See Supplementary Tables 3 and 5 for full domain annotations.

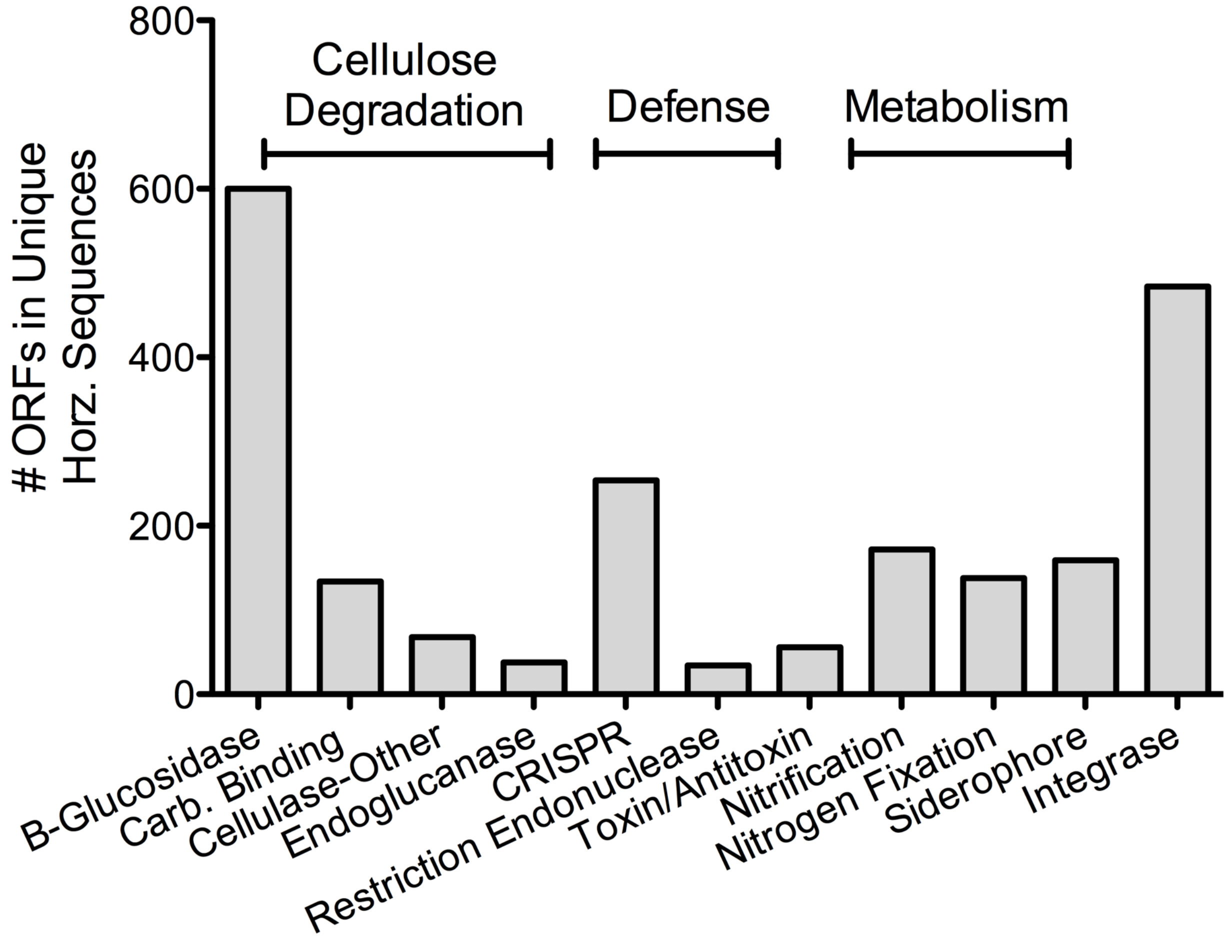


**Supplementary Figure 5: Ecologically significant genes.**

Assembly of DNA sequence reads uniquely found in horizontal communities at time point $T_1$, followed by interrogation of the Conserved Domain Database (CDD) identified predicted proteins with likely ecological relevance (see Extended Data Table 4 for domain descriptions and abundances).

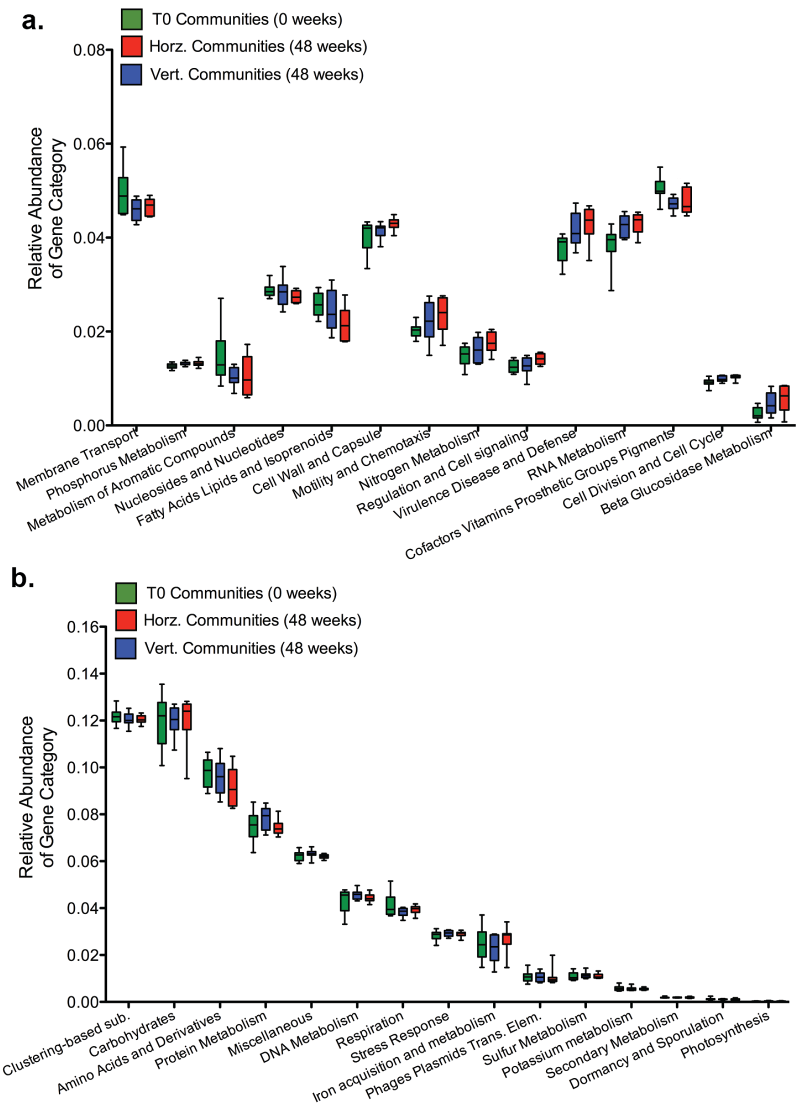


**Supplementary Figure 6: Functional gene categories.**

**a,** Relative abundances of specific gene categories that showed elevated differences by visual inspection of the data in either vertical or horizontal communities compared to respective $T_0$ founding communities. **b,** Relative abundances of specific gene categories that were highly similar in either vertical or horizontal communities compared to respective $T_0$ founding communities. Data are shown as box and whisker plots depicting median, inter quartile range (box) and full data spread from 10 replicate mesocosms.

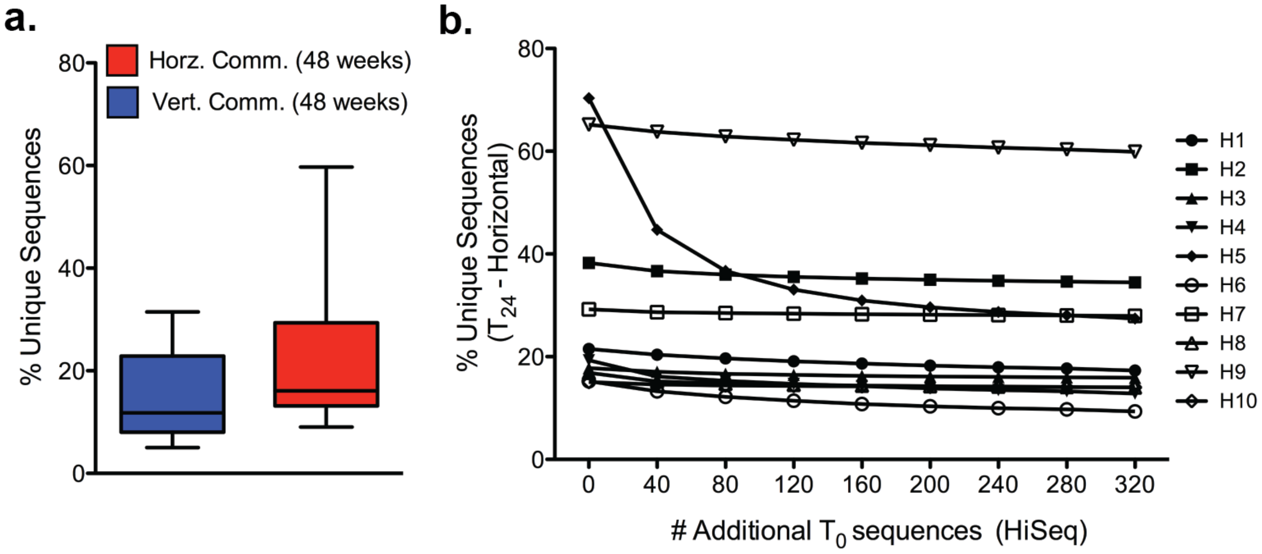


**Supplementary Figure 7: Distinguishing sequences transferred by SGEs from rare sequences.**

**a,** Percentage of sequences found at $T_{24}$ (48 weeks), but not present at $T_0$, thus identified as "unique" sequences. Data are shown as box and whisker plots depicting median, inter quartile range (box) and full data spread from 10 replicate communities. **b,** To determine the impact of increasing sequencing depth, $T_0$ communities were sequenced on the HiSeq platform delivering ~320 million sequence reads per community. Each was then arbitrarily divided into eight 40 million sequence read metagenome data sets, and the "unique" sequence analysis pipeline applied in order to identify any new matching sequences (see Supplementary Methods Fig. 1). Newly discovered sequences where then removed and the "unique" analysis pipeline re-applied after addition of the second $T_0$ metagenome. The process was repeated with all eight $T_0$ metagenomes showing the impact of additional sequencing on discovery of unique sequences.

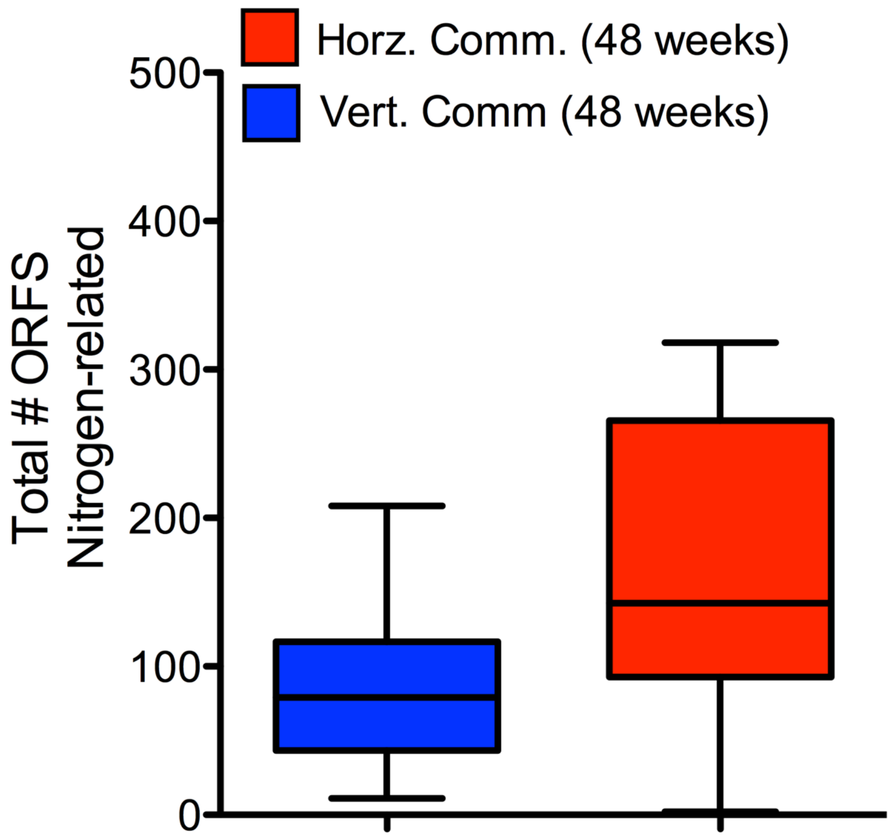


**Supplementary Figure 8: Nitrogen-related genes are enriched in "unique" horizontal sequences.**

A total of 136 nitrogen related domains were extracted from the Conserved Domain Database (CDD) based on their presence in at least one of the twenty "unique" assemblies. The abundance of all domains was determined. Data are shown as box and whisker plots depicting median, inter quartile range (box) and full data spread from 10 replicate communities. See Supplementary Table 9 for domain descriptions and abundance values.

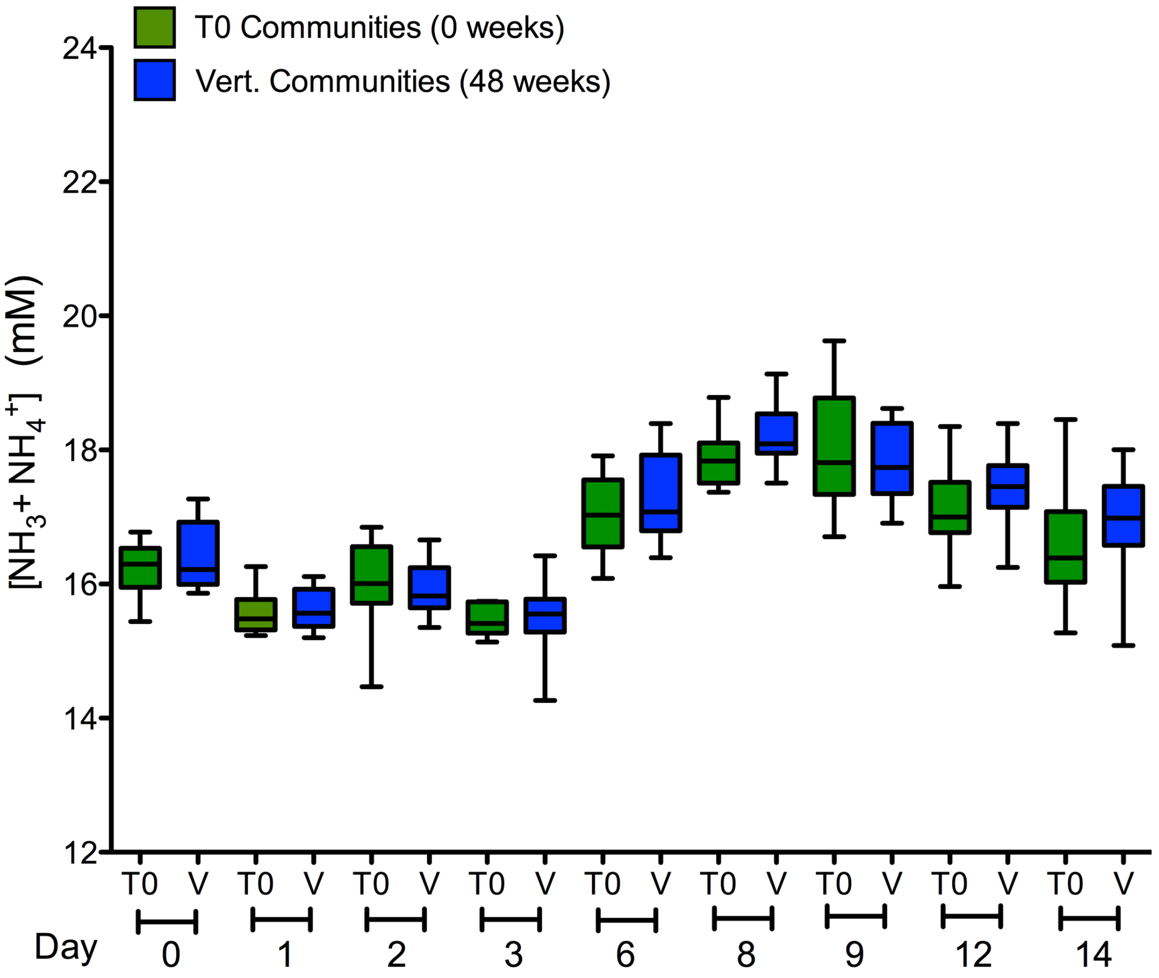


**Supplementary Figure 9: No change in function of communities subject to the vertical transfer regime after 48 weeks.**

Ammonia / ammonium concentration on selected days of the two-week culture period re-founded by $T_0$ and in $T_{24}$ communities subject to the vertical transfer regime. Data are shown as box and whisker plots depicting median, inter quartile range (box) and full data spread from 10 replicate communities (triplicate measures were obtained from each community on each occasion).

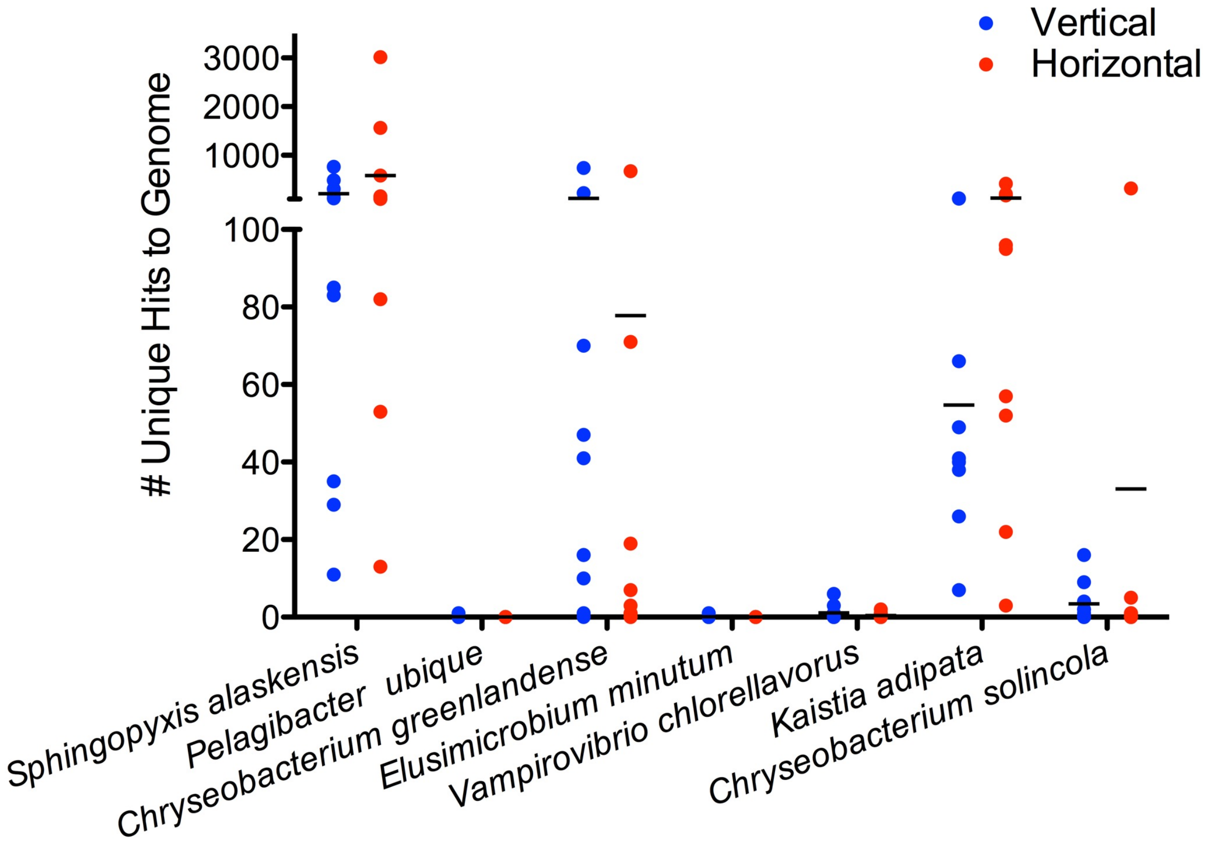


**Supplementary Figure 10: DNA sequence reads mapping to ultra-microbacteria.**

Recognising that a class of minuscule bacteria — the ultra-microbacteria — could potentially escape sedimentation and pass through a 0.22 µm filter DNA sequence reads from $T_{24}$ horizontal communities and vertical communities were mapped to the available reference genomes.

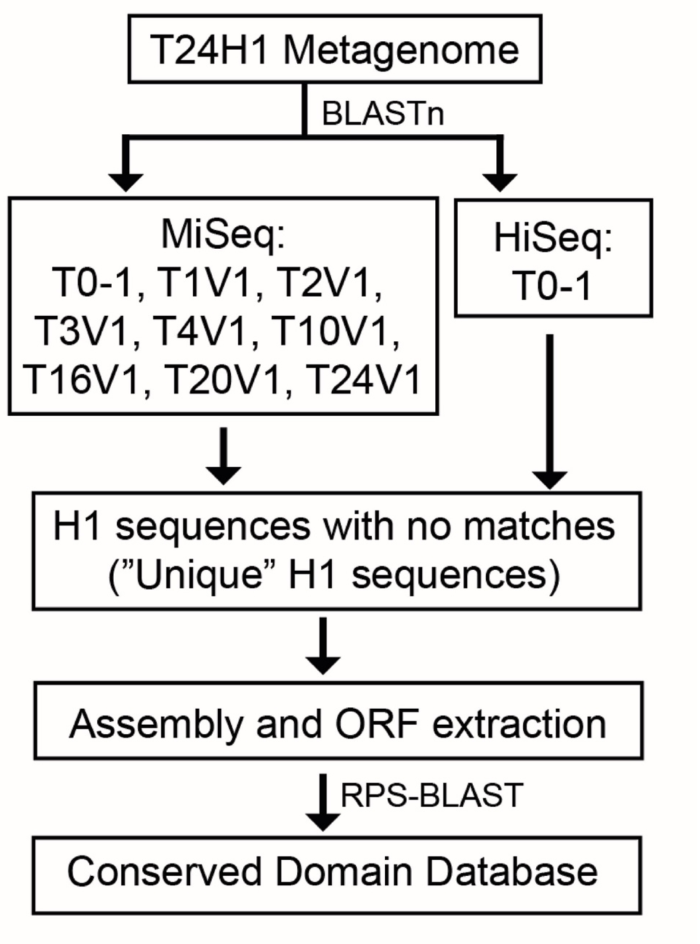


**Supplementary Methods Figure 1: Pipeline for identifying "unique" sequences.**

Metagenomes from horizontal communities were compared to their paired vertical community metagenomes at eight separate eight time points ($T_1$, $T_2$, $T_3$, $T_4$, $T_{10}$, $T_{16}$, $T_{20}$, $T_{24}$) as well as the metagenome of their founding $T_0$ community using BLASTn. Time point and initial $T_0$ metagenomes were sequenced using the MiSeq platform. To increase the depth of coverage the founding $T_0$ communities were also sequenced using the HiSeq platform. Horizontal sequences that matched Vertical or $T_0$ metagenomes were removed, leaving a set of sequences that were "unique" to each HC and were absent (undetectable) in any of the paired VCs or founding $T_0$ communities. "Unique" horizontal sequences were assembled, Open Reading Frames (ORFs) extracted, and analyzed using the Conserved Domain Database (CDD). As a control the same analysis was performed using the $T_{24}$ metagenomes from vertical communities to identify "unique" vertical sequences.
